# Dynamic Reprogramming of Stromal Pdgfra-expressing cells during WNT-Mediated Transformation of the Intestinal Epithelium

**DOI:** 10.1101/2025.01.22.634326

**Authors:** O Pellon-Cardenas, P Rout, S Hassan, E Fokas, He Ping, I Patel, J Patel, O Plotsker, A Wu, R Kumar, M Akther, A Logerfo, S Wu, DE Wagner, D Boffelli, KD Walton, E Manieri, K Tong, JR Spence, NJ Bessman, RA Shivdasani, MP Verzi

## Abstract

Stromal fibroblasts regulate critical signaling gradients along the intestinal crypt-villus axis^1^ and provide a niche that supports adjacent epithelial stem cells. Here we report that *Pdgfra*-expressing fibroblasts secrete ligands that promote a regenerative-like state in the intestinal mucosa during early WNT-mediated tumorigenesis. Using a mouse model of WNT-driven oncogenesis and single-cell RNA sequencing (RNA-seq) of mesenchyme cell populations, we revealed a dynamic reprogramming of Pdgfra^+^ fibroblasts that facilitates WNT-mediated tissue transformation. Functional assays of potential mediators of cell-to-cell communication between these fibroblasts and the oncogenic epithelium revealed that TGFB signaling is notably induced in Pdgfra^+^ fibroblasts in the presence of oncogenic epithelium, and TGFB was essential to sustain regenerative-like growth of organoids *ex vivo*. Genetic reduction of *Cdx2* in the β-catenin mutant epithelium elevated the fetal-like/regenerative transcriptome and accelerated WNT-dependent onset of oncogenic transformation of the tissue *in vivo*. These results demonstrate that Pdgfra^+^ fibroblasts are activated during WNT-driven oncogenesis to promote a regenerative state in the epithelium that precedes and facilitates formation of tumors.

## Introduction

Colorectal cancer (CRC) is a leading cause of cancer-related deaths worldwide.^2^ Morphogenesis and homeostasis of the adult mammalian intestine require strict tuning of the Wnt/β-catenin pathway because elevated WNT signaling drives the initiation and progression of most human and mouse intestinal cancers.^3–8^ Canonical WNT signals stabilize the dual signaling/adhesion protein β-catenin. In the absence of a WNT ligand, cytosolic β-catenin is targeted for proteosome degradation by a dynamic multiprotein destruction complex. WNT signaling inhibits beta-catenin proteolysis, followed by its translocation and accumulation in the cell nucleus, where it transactivates target genes.^9,10^ While signaling changes in the epithelium have been studied in depth, any resulting changes in the underlying stroma during oncogenesis are less clear, owing to the complexity of sub-epithelial cellular heterogeneity. Recent studies begin to uncover details about how stroma-derived ligands, such as prostaglandins, might support tumor-initiating epithelial stem cells.^11^

Fibroblasts preserve organ architecture and can exhibit activated states in response to tissue perturbations.^12^ Across tumors of different origin, fibroblasts give rise to specialized cancer-associated fibroblasts (CAFs) with antigen-presenting (apCAFs), inflammatory (iCAFs), or myofibroblastic (myCAFs) properties, each with different outcomes on tumorigenesis.^13^ Intestinal fibroblasts also execute myriad functions that reflect their complex heterogeneity, such as determination of epithelial cell fates and priming of immune cells.^14^ Mesenchymal fibroblasts are a critical source of Wnts, among other intestinal stem cell (ISC) niche factors that maintain intestinal crypt homeostasis.^1,15–21^ The largest subset, which expresses *Pdgfra*, impacts decisions on proliferation versus differentiation of ISCs by generating a BMP gradient along the stem and differentiated cell compartments.^1,22^. Fibroblasts expressing high or low *Pdgfra* have unique transcriptional and functional profiles in homeostasis and diseased tissue.^1,11,23^ Based on their significant role in tissue physiology, it is important to understand the function of these cells in tumorigenesis.

In response to various forms of epithelial damage, a gene signature that was first described for its enrichment in fetal intestinal organoid cultures marks regenerative crypt epithelial cells.^24–28^ In this regenerative state, dedifferentiation of *Lgr5-*negative cells is sustained by a YAP/TAZ gene signature that coincides with markers of late fetal intestinal organoids.^29^ Transcriptional reprogramming to this regenerative state is also invoked during initiation and spread of colorectal tumors.^30,31^ In advanced stages, metastatic tumor cells carry the most primitive undifferentiated state, associated with loss of intestinal identity.^31^ Accumulation of oncogenic mutations in tumor cells can drive WNT-independent growth by TGFB-mediated “lineage reversion”.^32^ Accordingly, human CRCs of the CMS4 type, characterized by high stromal infiltration and TGFB activation^33^, harbor a large population of *Clusterin*-expressing cells with a YAP-activated signature attributed to ISC revival (revSCs).^34–36^ Screening of patient-derived organoid (PDOs) co-cultures with stromal and fibroblast cells has shown that CAF signaling promotes this revival state to induce cancer chemoresistance^37^.

Here we report that conditional activation of WNT signaling via β-catenin gain-of-function promptly induces a revival-like transcriptional state accompanied by a YAP-associated gene signature. This oncogenic state appears to be supported by regenerative ligands produced by *Pdgfra*-hi and *Pdgfra*-lo fibroblasts in response to the onset of β-catenin-mediated hypertrophy. In enteroid-fibroblast co-cultures, elevated expression of these regenerative markers depends on paracrine TGFB signaling within the fibroblasts. Pdgfra-lo cells secreted pro-regenerative ligands in two different mouse models of WNT-driven intestinal tumors. Finally, compound mouse mutants reveal that partial loss of CDX2, which elevates the fetal/regenerative program, accelerates the oncogenic effects of mutant *β-catenin*.

## RESULTS

### Delayed transformation and initial preservation of differentiation ability in cells with oncogenic mutations in the WNT pathway

A notable delay separates the onset of oncogenic *Ctnnb1* mutation and the consequent transformation of epithelial morphology. Monitoring this transformation over time in *β-catenin^GOF/+^;Villin-Cre^ERT^*^2^ mice^7,38^ (Fig. 1A), we found that morphology typical of pre-cancerous lesions did not appear until approximately 14 days after expression of mutant *β-catenin^GOF^*. At this 2-week time point, crypt cells displayed a hematoxylin-rich appearance and increased cellular density compared to control animals. Over the next 2 weeks, this cellular morphology, previously described as a “crypt progenitor-cell” (CPC) phenotype^39^, expanded into the region normally occupied by intestinal villi until by 28 days post-induction of the *β-catenin^GOF^*mutation, when almost all of the duodenal mucosa was populated by crypt-progenitor-like epithelium (Fig. 1A, S1A). Epithelial transformation was most pronounced in the duodenum and decreased progressively towards the colon (Fig. S1B-C), though the delay in transformation was also observed in the colon, using a colon-specific Cre transgene (Fig S1B). This delay suggested that *β-catenin^GOF^* mutant cells differentiate normally over at least the first 2 weeks;. alternatively, mutant cells might loiter in the crypts without differentiation, allowing the CPC zone to expand gradually. To resolve these possibilities, we initiated the *β-catenin^GOF^* mutation in Lgr5+ intestinal stem cells using the *Lgr5-Cre^ERT^*^2^*-GFP* allele^40^ and followed mutant cells using the *ROSA-LSL-tdTomato* lineage tracing allele.^41^ Five to 7 days post-tamoxifen treatment, tdTomato-labeled cells appeared on the villi in both control and *β-catenin^GOF/+^* mutant intestines, with no difference between genotypes in the distance that labeled cells had traversed (Fig 1B-C). These data suggest that ISCs harboring a single *β-catenin^GOF^*mutant allele retain normal differentiation and migration rates.

**Figure 1.**
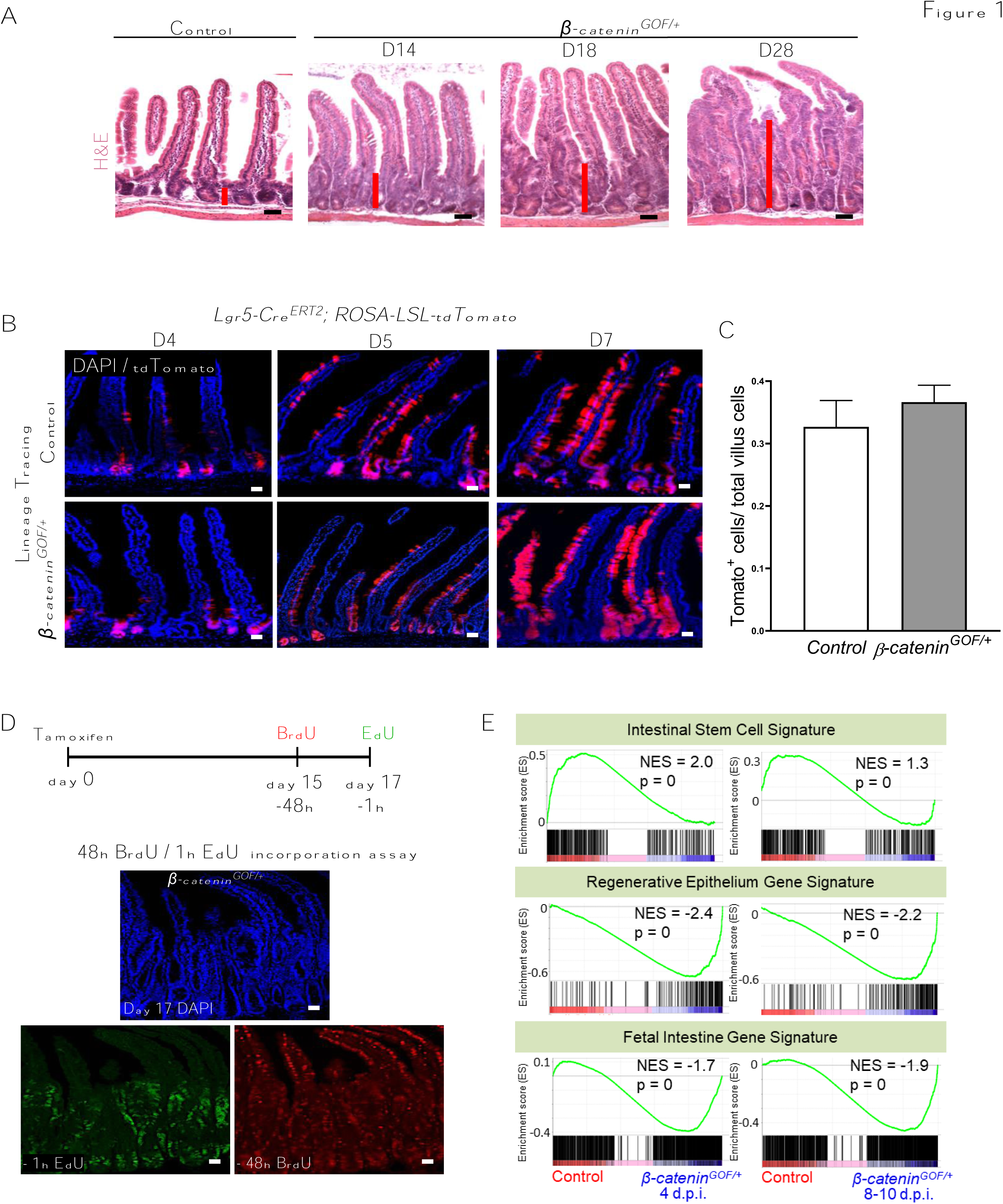
There is a delay in tissue transformation following oncogenic mutations activating the WNT pathway. (A) Histology (H&E) depicts a latency in tissue transformation upon WNT signaling activation. Red bar indicates degree of transformation as the expansion of crypt-progenitor zone with increasing days of activation of single mutant *β-catenin* allele in the proximal duodenum using the *Villin-Cre^ERT^*^2^ driver. (B) Lineage tracing of *β-catenin* mutant stem cells in the proximal duodenum of control or *β-catenin^GOF/+^; Lgr5-cre^ERT^*^2^ mice for 7 days reveals unperturbed luminal migration rates despite an elevated level of WNT signaling (n = 5 per group). (C) Quantitation of traced Td-tomato^+^ Lgr5^+^ cells in villus compartment per genotype at day 5 after expression of a *β-catenin^GOF^* mutant allele using the *Lgr5-Cre^ERT^*^2^ transgene (n = 5 per group). (D) Dual pulse labeling of *β-catenin* mutant stem cells with BrdU and EdU on day 17 of WNT-mediated tissue transformation show the continuing capacity of these mutant cells to differentiate. *β-catenin^GOF/+^; Villin-cre^ERT^*^2^ mice mice were injected with BrdU (48h pre-harvest) and EdU (1 h pre-harvest) to identify differentiated (red) and actively proliferating (green) cells (n = 3 biological replicates). (E) RNA-seq analysis of primary crypt epithelium shows activation of regenerative and fetal-like gene signatures in control vs *β-catenin^GOF/+^* crypt cells at day 4, or day 8-10 post-induction of a single mutant *β-catenin* allele. *Scale bars: 50 µm.

To determine whether cells in the expanded CPC zone 15 days post-treatment with tamoxifen could still differentiate and transit into occupy overtly normal villus epithelium, we injected mice with BrdU on day 15 and euthanized them 2 days later, thus marking cells that were in S-phase 48 h earlier. Additional EdU labeling 1 h before euthanasia marked cells actively cycling at the time of tissue harvest. Cells exhibiting differentiated villus morphology 17 days post-induction of the *β-catenin^GOF^* allele lacked EdU labeling (green), indicating that they were post-mitotic, but they carried the BrdU label, indicating that they were proliferative 2 days earlier (Fig. 1D). Consistent with a differentiated state, these BrdU+/EdU-cellls expressed the differentiation marker Alkaline Phosphatase (Fig. S1D). To investigate the molecular changes associated with *β-catenin^GOF^*activation, we examined bulk transcriptomes of epithelium isolated 4 days and 8-10 days after tamoxifen treatment. Increased *β*-catenin activity was associated with decreased markers of Lgr5+ ISCs and elevated expression of genes associated with epithelial regeneration following intestinal damage and with fetal versus adult intestinal organoids^28,42^ (Fig. 1E). A YAP signaling signature, a known mediator of intestinal regeneration and cancer progression,^29^ was also induced. In contrast, an intestine-specific WNT signaling signature^43^ lessened by day 8 after induction of the *β-catenin^GOF^* allele (Fig. S1E).

Taken together, these data indicate that *β-catenin^GOF^* mutant cells with a transformed morphology can still transition into a post-mitotic state with overt features of differentiation. Even after acquiring the mutation, ISCs and cells with the CPC phenotype can differentiate normally. Moreover, epithelial cells with elevated *β*-*catenin* signaling express transcripts characteristic of ISC regeneration after epithelial injury. The delay between Wnt activation and tissue transformation additionally suggests that other signals, potentially emanating from the stroma, contribute to the transformation.

### Stromal fibroblasts express pro-regenerative ligands in response to epithelial oncogene activation

Single-cell technologies have made diverse intestinal mesenchyme cell populations accessible to study^1,23,44^. To appreciate changes in non-epithelial cells in response to epithelial WNT activation, we conducted scRNA-seq early during oncogenic transformation. In *β-catenin^GOF/GOF^;Villin-Cre^ERT^*^2^ mice, tissue transformation and exacerbated cell proliferation were evident within 5 days of tamoxifen treatment (Fig. 2A-B). We collected intestines from control and experimental mice at different times post-treatment, removed smooth muscle and epithelial cells, and dissociated the remaining stroma into single cells for scRNA-seq (Fig. 2B), following methods described previously.^1^ All stromal cell constituents were detected as expected,^1^ including robust populations of immune and endothelial cells and fibroblasts (Fig. 2C-D). Cells from each isolation exhibited similar scRNA-seq quality metrics (Fig. S2A). On the background of a merged uniform manifold approximation and projection (UMAP – Fig. 2E), fibroblasts marked by *Pdgfra* expression were distributed asymmetrically, with control cells populating the right edge of the merged single-cell cluster and those isolated from the *β-catenin^GOF^* mice populating the left edge. This observation implies that epithelial *β-catenin* activation induces transcriptional changes in underlying fibroblasts (Fig. 2E-F, S2B-C). Pdgfra-hi sub-epithelial myofibroblasts, also called telocytes, line the epithelium with long cytoplasmic projections, while Pdgfra-lo cells sit deeper in the stroma and have been stratified into CD81+ and CD81-populations^1^; CD81+ trophocytes support epithelial growth *in vivo* and *in vitro*.^1^ Comparing these Pdgfra+ clusters with public datasets of fibroblast classification revealed transcripts associated with fibrotic and inflammatory cancer-associated fibroblasts^45,46^ in both Pdgfra cell clusters derived from *β-catenin^GOF^* mice (Fig S2D). These transcripts increased over time after *β-catenin^GOF^* activation (Fig. S2D). The sum of these findings prompted us to investigate whether stromal cells, and which subpopulation, might influence the speed of oncogenic transformation.

**Figure 2.**
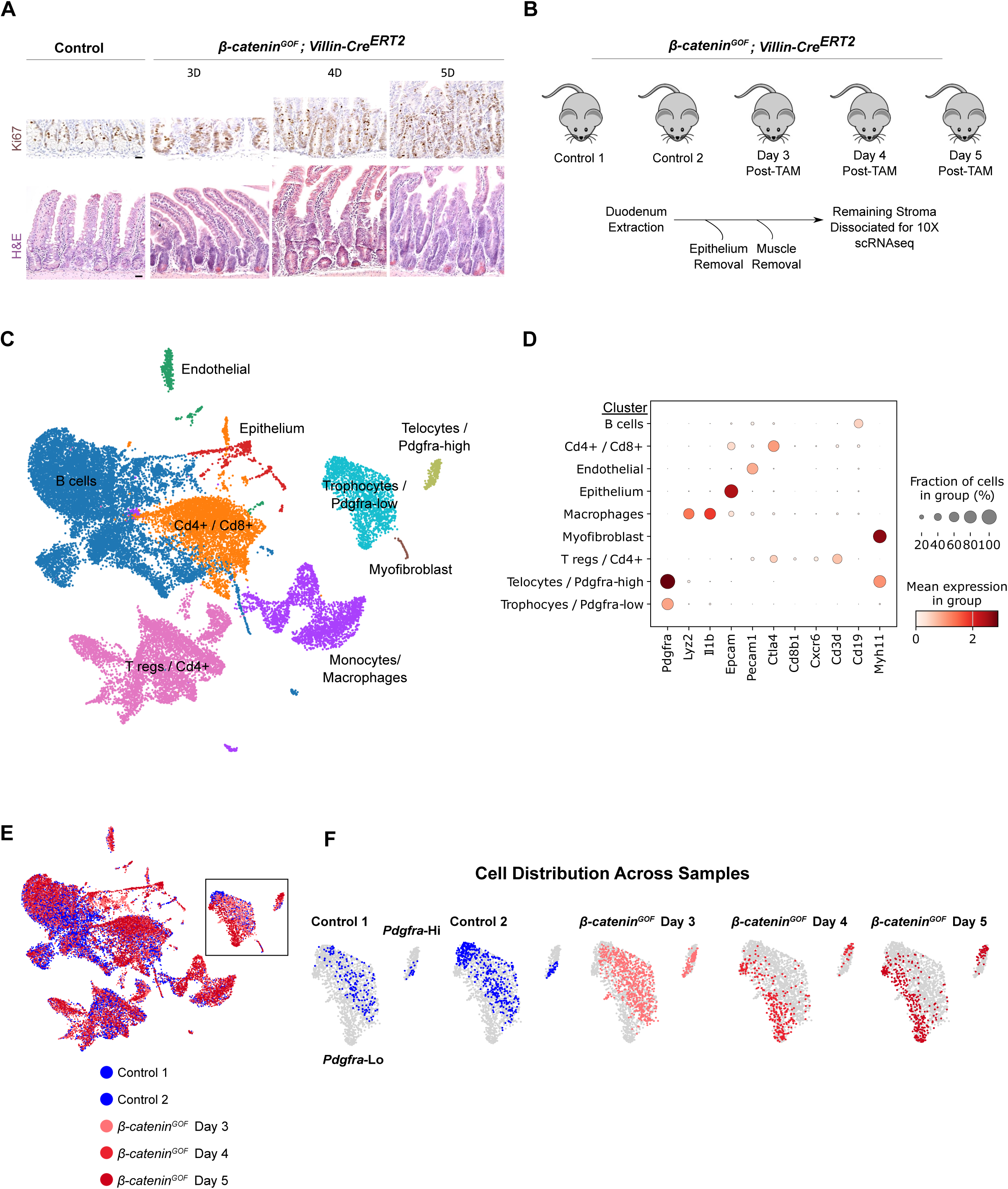
Survey of mesenchymal cell populations during WNT-mediated tissue transformation reveals a transcriptomic shift in Pdgfra+ fibroblasts. (A) Immunohistochemistry (IHC) of Ki67 and Histology (H&E) depicts rapid onset of jejunum tissue transformation across a time course of homozygous activation of mutant *β-catenin* using the *Villin-Cre^ERT^*^2^ transgene. Duodenum samples were used for scRNA-seq (n = 3 biological replicates). Scale bars: 50 µm. (B) Experimental schematic for the isolation of mesenchymal cells during WNT-mediated tissue transformation. (C) UMAP depicts cell populations of the intestinal stroma. UMAP is a composite of all cells from 5 experimental samples (2 control and 3 mutants). (D) Dotplot indicates expected marker gene expression to identify mesenchyme cell types. (E-F) UMAP depicts an asymmetric distribution, particularly in Pdgfra-lo and Pdgfra-hi mesenchyme clusters (enlarged in F), indicating a transcriptomic shift in response to epithelial activation of *β-catenin^GOF^*.

### Pdgfra-lo fibroblasts direct transcriptional reprogramming of oncogenic epithelium

Primary cultures of Pdgfra-lo cells^1^ mirrored the properties of their *in vivo* counterparts, including GFP expression when cultured from *Pdgfra-H2B-eGFP* mice and the ability to support growth of spherical organoids without the addition of growth factors typically required to support enteroid monocultures^47^ (Fig. 3A – B). Co-culture of these Pdgfra-lo cells with control or *β-catenin^GOF^*enteroids induced spherical morphology and rapid expansion in size (Fig 3B). Enteroids growing in routine conditions that mimic homeostasis^47^ grow as branched structures, while spheroid growth is more typical of high-WNT conditions and regenerative states.^48,49^ In these co-cultures, we therefore examined expression of transcripts associated with homeostatic or restorative states. Regenerative markers *Clu* and *Anxa1* were consistently elevated in both control and *β-catenin^GOF^* enteroids under co-culture, while markers of normal crypt based columnar (CBC) cells were significantly diminished (Fig. 3C, Fig. S3A). Concordant with our *in vivo* findings (Fig. 1E), basal expression of regenerative markers was elevated in *β-catenin^GOF^*enteroid monocultures (Fig. 3C). Given that Pdgfra-lo fibroblasts appeared to sustain “regeneration-like” growth of both wild type and *β-catenin^GOF^* enteroids, we measured expression changes in potential ligands that mediate homeostatic vs regenerative states. WNT2B, GREM1, and RSPO3 are such ligands, responsible for proper localization of normal Lgr5^+^ ISCs in the crypt base^50,51^. Transcripts for these pro-ISC ligands were downregulated in both control and *β-catenin^GOF^*enteroid co-cultures (Fig. 3D). Among potential ligands that mediate ISC regeneration after injury,^11,34,52,53^ *Ereg*, *Il11*, *Ptgs2,* and *Tgfb1* were all elevated in Pdgfra+ mesenchymal cells following exposure to *β-catenin^GOF^* epithelium (Fig. 3E). Sequencing of RNAs in FACS-sorted *Pdgfra-H2B-GFP* expressing cells from the co-cultures (Fig S3B) confirmed that *Ereg*, *Tgfb1*, *Il11*, and *Ptgs2* were indeed elevated in wild type Pdgfra+ mesenchymal cells co-cultured with *β-catenin^GOF^*enteroids (Fig. 3F). Primary unfractionated mesenchymal cells freshly isolated from *β-catenin^GOF^* mutant intestines confirmed induction of these pro-regenerative ligands and independent whole-thickness intestinal samples confirmed induction of *Ereg* and *Il11* (Fig. 3G-H). Thus, oncogenic epithelium reproducibly induced pro-regenerative characteristics in Pdgfra-lo fibroblasts.

**Figure 3.**
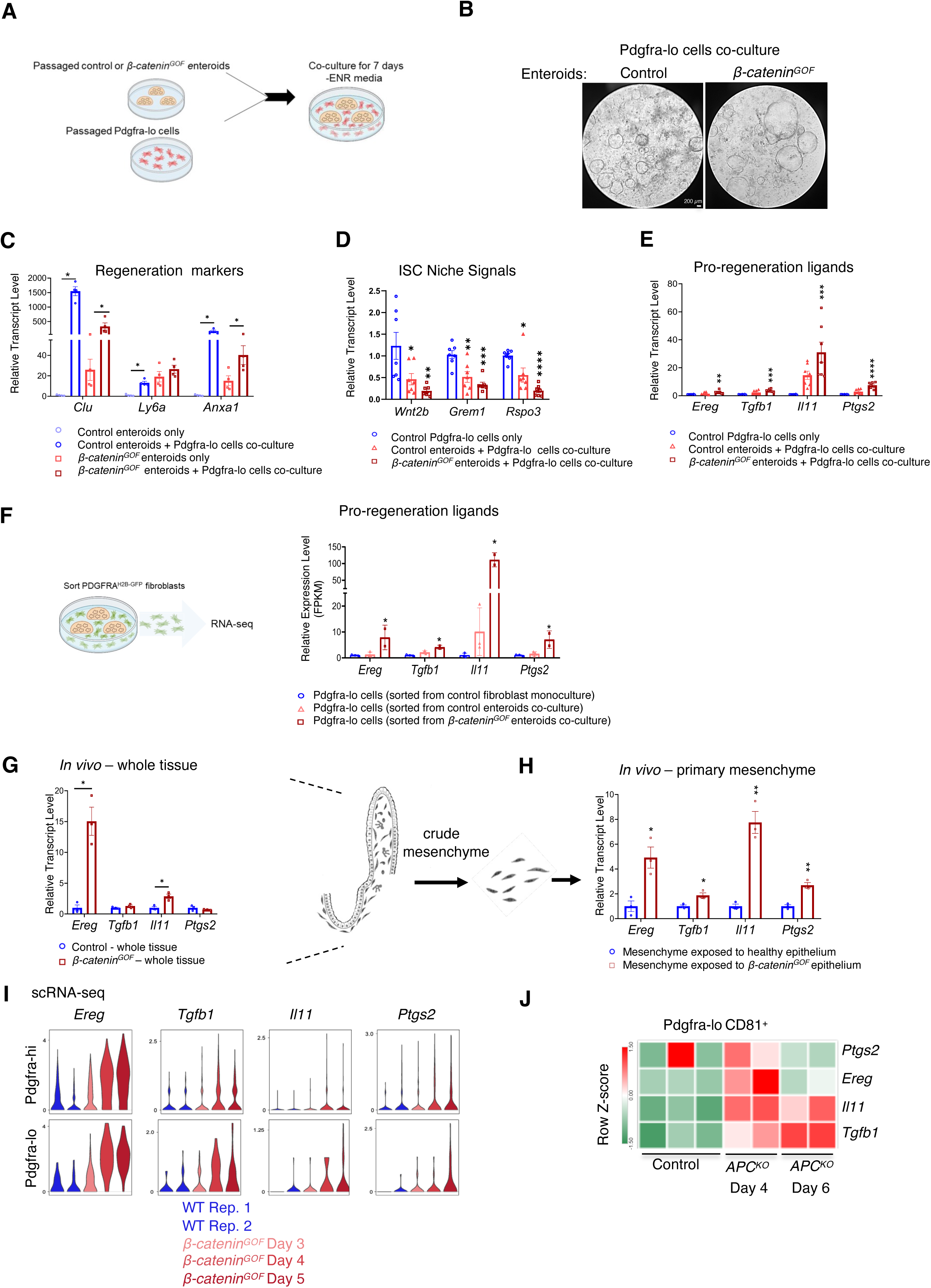
Stromal Pdgfra+ cells are a key source of ligands known to promote epithelial regeneration. (A) Schematic of co-culture experiments. Control or *β-catenin^GOF^* enteroids at passage 1 were co-plated with passaged Pdgfra-lo cells in Matrigel droplets for 7 days in –ENR media. (B) Pdgfra-lo cells promote hyperproliferative growth of control and *β-catenin^GOF^* enteroids that resemble fetal enteroids instead of branched structures. Scale bar = 200µm. (C) qRT-PCR analysis shows increased expression of epithelial regeneration markers in control and *β-catenin^GOF^ enteroids* grown in co-culture with Pdgfra-lo cells (n = 4 biological replicates, Mann– Whitney U-test, p < 0.05*). (D) ISC niche signals are downregulated in Pdgfra-lo cells co-cultured with control or *β-catenin^GOF^*enteroids (n = 7 biological replicates, one way ANOVA with Tukey post hoc test, p < 0.0001****, p < 0.001***, p < 0.01** and p < 0.05*). (E) Pro-regeneration ligands are elevated in Pdgfra-lo cells co-cultured with *β-catenin^GOF^* enteroids. (n = 7 biological replicates, one way ANOVA with Tukey post hoc test, p < 0.05*). (F) RNA-seq analysis of fluorescence-activated cell sorting (FACS) of Pdgfra-H2B-GFPcells confirm elevated expression of *Ereg, Tgfβ-1, Il11* and *Ptgs2* transcripts in Pdgfra-lo cells co-cultured with *β-catenin^GOF^* enteroids over 7 days (n = 3 biological replicates for Pdgfra-lo monocultures, n = 3 control enteroid-cocultures and n = 2 biological replicates for *β-catenin^GOF^* co-cultures, one way ANOVA with Tukey post hoc test, p < 0.001***, and p < 0.01**). (G) qRT-PCR data indicating upregulation of *Ereg and Il11* in whole tissue samples derived from *β-catenin^GOF^* mice after 4 days of tamoxifen injection (n = 3 biological replicates, Mann–Whitney U-test, p < 0.01** and p < 0.05*). (H) qRT-PCR of primary unfractionated mesenchyme isolated from *β-catenin^GOF^* mice show increased expression of pro-regeneration ligands (n = 3 biological replicates, Mann–Whitney U-test, p < 0.01** and p < 0.05*). (I) Violin plots showing increased expression levels per single cell for *Ereg, Tgfβ-1, Il11* and *Ptgs2* in Pdgfra-hi and Pdgfra-lo cells in a time course of β-catenin activation using *β-catenin^GOF^* mice. (J) Heatmap indicate sustained activation of *Tgfβ1* and *Il11* identified by RNA-seq analysis of Pdgfra-lo CD81^+^ cells isolated from *Apc^KO^* mice.

Pdgfra-lo cells derived from *β-catenin^GOF/+^* mutant mice were also consistently less efficient in supporting growth of wild-type enteroids than Pdgfra-lo cells isolated from healthy controls (Fig. S3C). We considered that this suppression of wild-type organoid growth might reflect elevated expression of BMP ligands and negative WNT regulators after Pdgfra-lo fibroblasts are exposed to oncogenic epithelium (Fig. S3D). These pro-differentiation signals are likely tolerated better by cells harboring oncogenic mutations compared to wild-type stem cells. Indeed, high levels of BMP2, a pro-differentiation ligand,^54^ did not affect long term viability of *β-catenin^GOF^* enteroids, but caused wild-type enteroids to collapse (Fig. S3E); this difference could confer a competitive advantage for mutant over normal epithelium. Additionally, re-examination of our scRNA-seq data revealed induction of regeneration-associated transcripts in both *Pdgfra*-expressing stromal cell populations (Fig. 3I). Thus, although Pdgfra-hi cells are not readily amenable to co-culture assays, they do acquire a transcriptional shift similar to that of Pdgfra-lo cells in response to oncogenic epithelial exposure.

To validate Pdgfra-lo cell expression of regenerative genes in another genetic model of acute WNT activation, we examined their gene expression changes in response to depletion of *Apc^f/f^* (a conditional mutant ^55^) in the epithelium using the *Villin-Cre^ERT^*^2^ driver. As in the *β-catenin^GOF^* model of WNT hyperactivation, RNAseq analysis of Pdgfra-lo CD81^+^ cells sorted 4 days after *Apc* inactivation in epithelial cells showed induction of *Ereg*, *Tgfb1*, *Il11*, and *Ptgs2* transcripts, and on the 6^th^ day, *Il11* and *Tgfb1* remained elevated in these cells (Fig. 3J). Together, these findings reveal that Pdgfra-lo stromal cells respond rapidly to oncogene activation in the overlying epithelium by suppressing homeostatic growth and expressing genes that ordinarily promote ISC regeneration.

### *Epiregulin* is dispensable for WNT-driven tissue transformation

CellChat, an algorithm that uses curated literature to match receptor-ligand pairs between cell types in scRNA-seq datasets,^56^ imputed that the EREG-EGFR signaling pathway was notably active between Pdgfra+ and mutant *β-catenin^GOF^* epithelial cells compared to control epithelial cells. Signaling was predicted to occur mainly from Pdgfra+ cells towards the transformed epithelium, with additional autocrine and paracrine signaling between Pdgfra-hi and Pdgfra-lo cells (Fig. 4A). In a survey of known EGF-family ligands, *Epiregulin* (*Ereg*) was the most notably induced in Pdgfra+ cells in response to epithelial *β-catenin^GOF^*activation (Fig. 4B-C); the *Egfr* receptor, which enables EPIREGULIN signal transduction,^57^ was also elevated in Pdgfra+ cells exposed to WNT-transformed epithelium (Fig S4A-B). Fluorescent RNA in situ hybridization in control and *β-catenin^GOF^* mutant intestines gave modest *Ereg* signal in control intestines and notably increased signals in stromal and epithelial cells of mutant animals (Fig. 4D). To test the role of EREG in promoting enteroid growth and tumorigenesis, we co-cultured control or *β-catenin^GOF^* enteroids with control or *Ereg^null^* mesenchyme isolated from *Ereg* knockout mice.^58^ *Ereg^null^* Pdgfra+ cells did not impact the growth or morphology of co-cultured enteroids (Fig. 4SC) or change expression of regenerative genes (Fig. 4SD). Nor was the rate of tissue transformation or induction of regenerative markers altered in *β-catenin^GOF/+^*;*Ereg*^-/-^ mice versus *β-catenin^GOF/+^*; *Ereg*^+/+^ controls (Fig. 4E-G). Epithelial *Ereg* loss also did not result in overt defects, and *Ereg^null^* crypts yielded healthy branched enteroids with differentiation potential similar to control crypts (Fig. S4E-F). Addition of EREG to enteroid cultures devoid of EGF did not alter homeostatic growth (Fig. S4G) or affect regenerative growth of primary and passaged *β-catenin^GOF^* enteroids (Fig. S4H-I). Altogether, these findings identify EREG as a dispensable factor in early oncogenic, WNT-driven expansion of the crypt-progenitor zone in our model.

**Figure 4.**
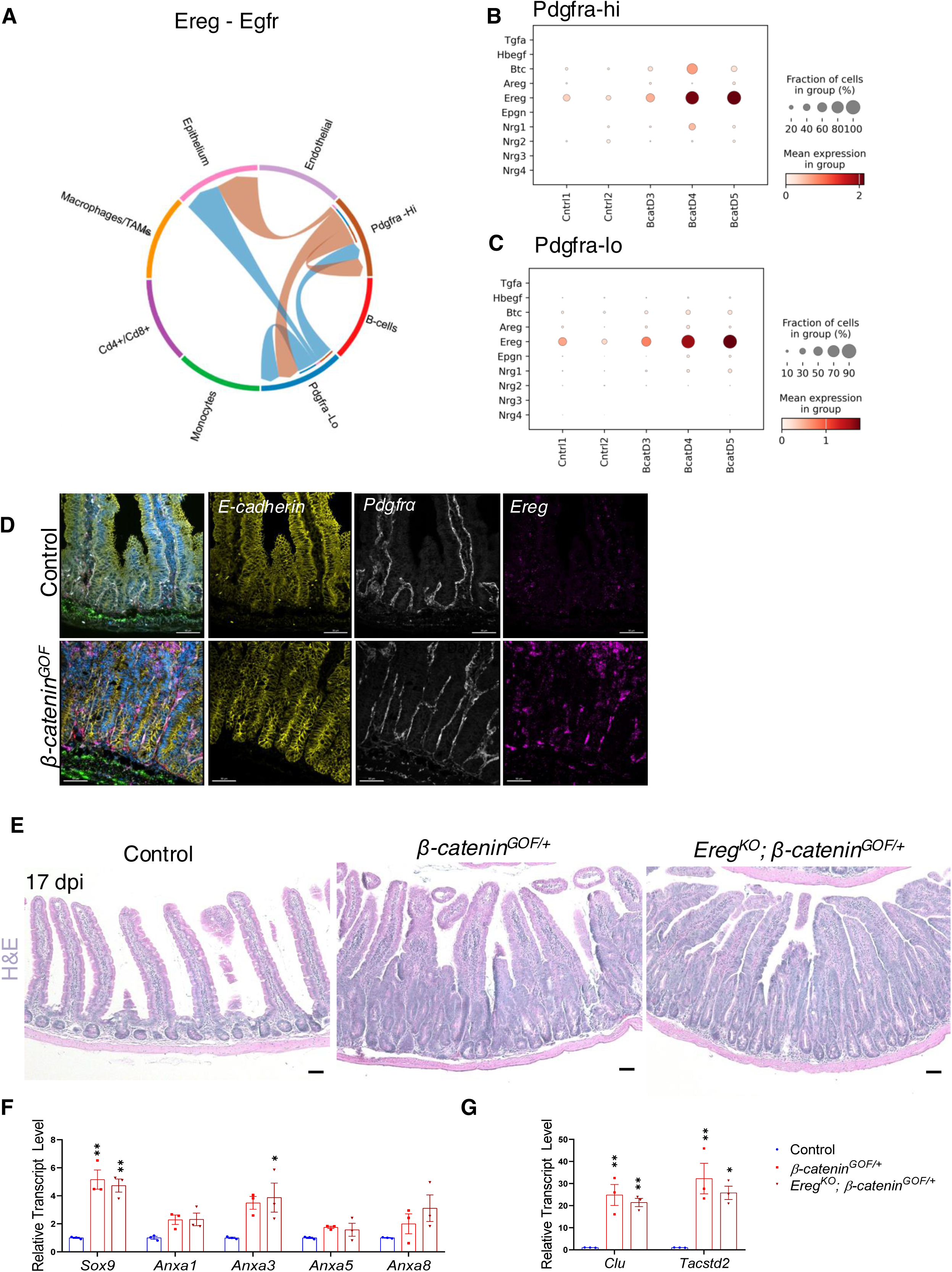
*Ereg* expression is induced in stromal fibroblasts in response to oncogenic change in the epithelium and is dispensable for tissue transformation. (A) Chord diagram derived from scRNA-seq data predicts that the Epiregulin ligand is utilized by Pdgfra+ mesenchyme cells to communicate with each other and the epithelium. (B-C) Dotplot of scRNA-seq data showing that *Ereg* expression is predominantly higher than other EGFR ligands in Pdgfra-hi and Pdgfra-lo cells across a time-course of *β-catenin^GOF^* activation. (D) RNAscope localized *Ereg* transcripts relative to Pdgfrα immunostaining and E-cadherin immunostaining (epithelial marker). Upregulation of *Ereg* is distinctively observed in Pdgfra cells and in certain regions of *β-catenin^GOF^* epithelium (n = 3 biological replicates). Scale bar = 50 µm. (E) Histology (H&E) of indicated mutant mouse duodenum shows that *Ereg* is dispensable to tissue transformation in the proximal duodenum of *β-catenin^GOF/+^* mice after 17 days of tamoxifen injection (n = 3 biological replicates). (F-G) qRT-PCR data indicate an equivalent induction of fetal-like intestine markers in transformed primary epithelium isolated from *β-catenin^GOF/+^; Villin-Cre^ERT^*^2^ and *Tgfbr2^KO^;β-catenin^GOF/+^; Villin-Cre^ERT^*^2^ mice (n = 3 biological replicates, one way ANOVA with Tukey post hoc test, p < 0.01** and p < 0.05*).

### Elevated TGFB signaling promotes regeneration-like behavior in early tumorigenesis

*Tgfbr2* and *Cald1*, genes related to stromal TGFB activation were among the top genes upregulated in both Pdgfra-Lo and Pdgfra-hi cells in response to sustained epithelium WNT activation (Fig. 5A, Fig. S5A), and *Tgfbr2* transcripts were particularly induced in Pdgfra+ cells compared to other stromal populations (Fig. 5A). TGFβ-1 drives the regenerative response to epithelial injury^27^ and may act on epithelial and mesenchyme cell populations^27^. To test for TGFB effects in stromal fibroblasts, we first defined a “TGFβ-1-induced stromal fibroblast gene signature” by treating cultures of primary *Pdgfra-lo-H2B-GFP* cells with exogenous TGFβ-1 and conducting RNA-seq (Fig. 5B-C, Fig. S5B). In parallel, we used *Tgfbr2^KO^* cells to measure any non-canonical TGFB1-induced activities, but few were observed (Fig. S5B). TGFβ-1 induced expression of regenerative ligands, such as *Wnt5a* and *Tgfb1* itself, while the BMP antagonist *Grem1,* which supports homeostatic stem cell activity,^50^ was repressed, indicating that *Pdgfra-lo-H2B-GFP* cells can be reprogrammed to support regenerative epithelial growth (Fig. S5C). We also observed activation of *Cald1* and *Igfbp7*, classic markers of TGFB activation in cancer associated fibroblasts^59^ (Fig. S5C). All these transcriptional changes reflect canonical TGFB signaling, as they were not observed in *Tgfbr2* knockout Pdgfra-lo-H2B-GFP cells (Fig. S5B-C). In our scRNA-seq data, Pdgfra-lo cells upregulated this newly defined TGFβ1-induced stromal fibroblast gene signature after β-catenin hyperactivation in the gut epithelium (Fig. 5C, Fig. S5A); among TGFβ family members, *Tgfb1* was notably induced (Fig. S5D). Genes upregulated in Pdgfra-lo cells exposed to oncogenic epithelium in vivo or those co-cultured with *β-catenin^GOF^*enteroids and then analyzed by bulk RNA-seq were also significantly correlated with the TGFB1-induced gene signature (Fig. 5D-E). These data show that TGFB activity is enhanced in Pdgfra+ cells following oncogenesis in vivo and can be recreated ex vivo. Thus, TGFB1 signaling may support both oncogenic transformation and the regenerative marker gene expression observed during early oncogenesis (Fig. 5C-E). To test these possibilities, we turned to enteroid monocultures and enteroid – Pdgfra-lo cell co-cultures.

**Figure 5.**
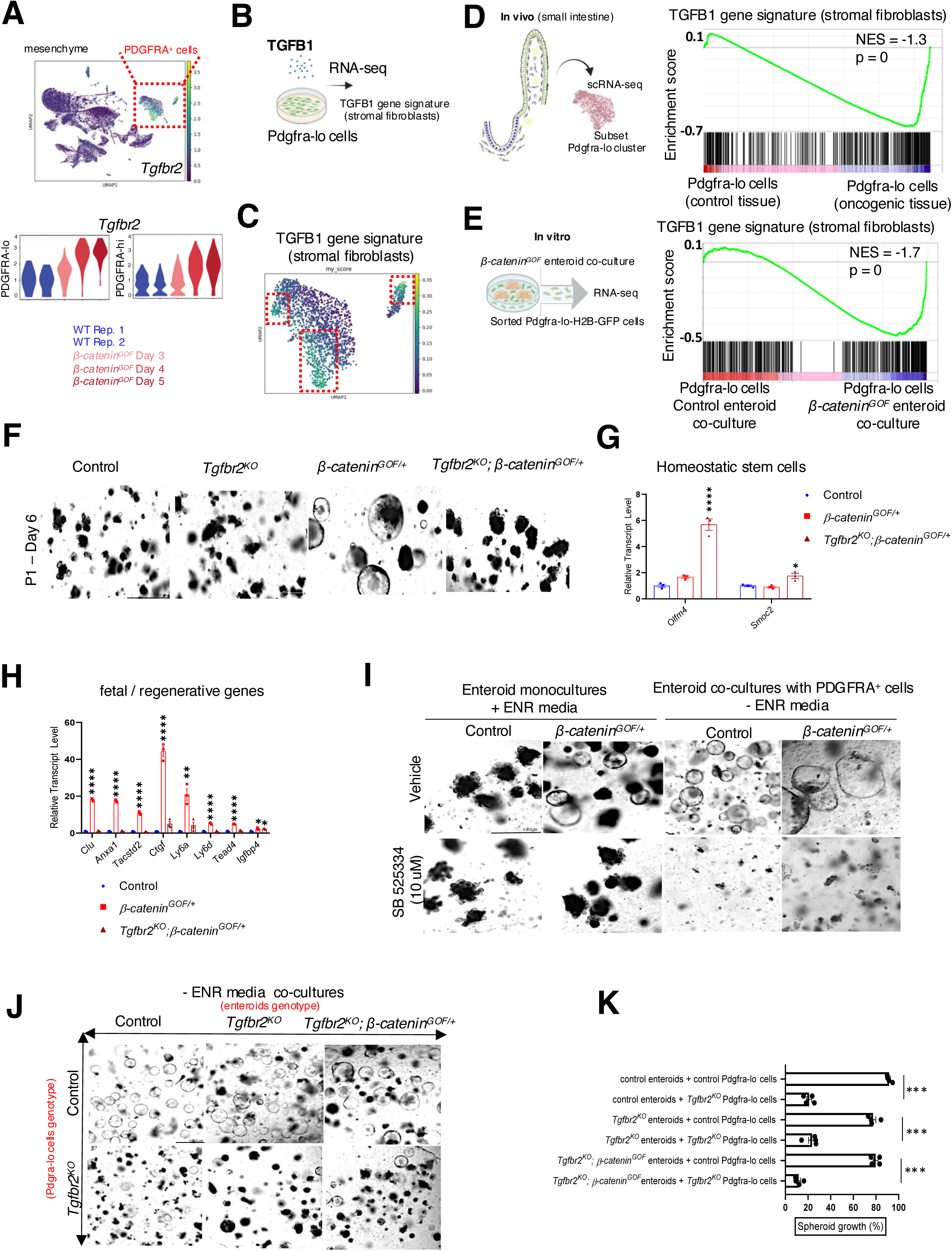
TGFB ligands induce stromal fibroblasts to express pro-regenerative ligands. (A) UMAP of all mesenchyme cell clusters indicates high *Tgfbr2* expression in *Pdgfra*-expressing cells. Violin plots indicate higher expression of *Tgfbr2* in both Pdgfra-hi and Pdgfra-lo cell populations after activation of *ß-catenin^GOF^* in the epithelium. (B) To define a TGFB1-gene signature (stromal fibroblasts), RNA-seq was conducted on Pdgfra-lo cells treated with either vehicle or recombinant TGFB1 (10 ng/mL) for 72 hours (also see Fig. S4A-B). (C) UMAP of Pdgfra mesenchyme cell clusters indicating an enrichment of a TGFB1 gene signature (stromal fibroblasts) in both Pdgfra-hi and Pdgfra-lo cell populations after epithelial *ß-catenin^GOF^* activation. (D-E) GSEA analysis showing enrichment of a TGFB1 gene signature (stromal fibroblasts) in Pdgfra-lo cells *in vivo* after exposure to oncogenic epithelium (D, differential gene expression analysis via scRNAseq), or *in vitro* (E with Pdgfra-lo cells sorted from co-cultures with *ß-catenin^GOF^* enteroids via bulk RNA-seq). (F) Depletion of epithelial *Tgfbr2* does not impact enteroid growth after passage and it prevents spheroid morphology normally induced by activation of the *ß-catenin^GOF^* mutant allele in enteroids. Primary organoids were derived from tamoxifen-injected mice and their growth was monitored up to six days of the first passage. (n = 3 biological replicates). Scale bar = 1000 µm. (G) qRT-PCR analysis of primary organoids at day 6 derived from control, *ß-catenin^GOF^; Villin-cre^ERT^*^2^ and *Tgfbr2^KO^*; *ß-catenin^GOF^; Villin-cre^ERT^*^2^ mice to test for stem cell markers (n = 3 biological replicates, one way ANOVA with Tukey post hoc test, p < 0.0001**** and p < 0.05*) (H) qRT-PCR data shows that fetal/regenerative reprogramming is inhibited in *Tgfbr2^KO^*; *ß-catenin^GOF^; Villin-cre^ERT^*^2^ mice (n = 3 biological replicates, one way ANOVA with Tukey post hoc test, p < 0.0001****, p < 0.01** and p < 0.05*) (I) Enteroid monocultures are viable after treatment with SB525334 TGFBR2 inhibitor (10 µM) over 5 days. However, susceptibility to SB525334 is observed in co-cultures of Pdgfra+ cells with control or *ß-catenin^GOF^* enteroids in –ENR media, indicating dependency of TGFB signaling in epithelial growth sustained by mesenchyme (n = 3 biological replicates). Scale bar = 1000 µm. (J-K) Genetic inactivation of *Tgfbr2* in enteroids does not prevent growth of enteroids in a co-culture setting. However, genetic inactivation of *Tgfbr2* in Pdgfra-lo cells significantly reduced spheroid growth of enteroids in co-cultures even when the *ß-catenin^GOF^* mutant allele is active in the enteroids (n = 4 biological replicates). Scale bar = 1000 µm.

Monoallelic *β-catenin^GOF^* mutation triggers spherical enteroid growth.^60^ Although TGFB signaling is dispensable to sustain homeostatic enteroid growth in standard ENR (EGF, NOGGIN, RSPONDIN1) cultures,^48^ *Tgfbr2* receptor deletion in enteroids reversed *β-catenin^GOF^*-induced spheroid growth (Fig. 5F), with *Tgfbr2^KO^;β-catenin^GOF/+^*enteroids taking on the branched morphology typical of homeostatic growth (Fig. 5F). Like *β-catenin^GOF/+^*controls, *Tgfbr2^KO^;β-catenin^GOF/+^* enteroids showed elevated expression of homeostatic stem cell markers, but did not activate regeneration markers (Fig. 5G-H). Thus, TGFB signaling is required to facilitate β-catenin-mediated regenerative reprogramming and, in the absence of mesenchyme, *β-catenin^GOF/+^* enteroids can autonomously invoke a regeneration-like state in a *Tgfbr2*-dependent manner (Fig 5H).

We further explored TGFB1 activity in the context of enteroid-fibroblast co-cultures, which occur in the absence of ENR factors. We co-cultured control and *β-catenin^GOF/+^* enteroids with Pdgfra-lo cells and used genetic and pharmacologic approaches to block TGFB signaling. Both control and *β-catenin^GOF/+^* enteroids co-cultured with Pdgfra-lo cells failed to grow when TGFBR was inhibited with 10 uM SB525334 (Fig. 5I). These TGFBR inhibitor concentrations did not affect growth in standard enteroid conditions,^48^ highlighting a specific dependency for TGFB1 signaling to mediate mesenchyme-supported epithelial growth (Fig. 5I). Genetic inactivation of *Tgfbr2* in Pdgfra-lo cells also hampered growth of co-cultured organoids, while enteroids lacking *Tgfbr2* were unaffected in co-cultures, further indicating a role for TGFB1 signaling in stromal support of epithelial expansion (Fig. 5J-K). Thus, TGFB1 not only induces a regenerative phenotype in enteroid cultures (Fig. S5E), it also exerts a large influence on Pdgfra+ cell reprogramming to promote regenerative growth. Indeed, treatment of Pdgfra-lo cells with TGFB1 induced expression of pro-regenerative ligands such as *Il11* and *Ptgs2* (Fig. S5F). Importantly, while treatment with TGFBR1 inhibitor blocked growth of enteroid co-cultures, it did not compromise Pdgfra-lo cell viability (Fig. S5G). Together, these experiments indicate that TGFB1 signaling, which is elevated during early oncogenesis, plays a critical role in stromal fibroblasts to support a regeneration-like state in the overlying epithelium.

### The fetal-associated regenerative program accelerates oncogenic transformation

In response to constitutive Wnt signaling by APC inactivation, intestinal epithelial cells were recently reported to express a fetal-like/regenerative program during the onset of colorectal cancer.^30^ Consistent with activation of a regenerative state in the *β-catenin^GOF^* model, we observed YAP accumulation in intestinal crypts and increased expression of SOX9 throughout the transformed epithelium (Fig. 6A). *β-catenin^GOF^* crypt epithelium also increased expression of regeneration markers (Fig. 6B). To test whether the fetal-like/regenerative state is important during early transformation, we reduced levels of CDX2, a transcription factor that acts as a barrier to that state; CDX2 loss from the adult epithelium correlates with activation of genes selectively elevated in fetal enteroids^49,61–63^. To reduce CDX2 levels in the context of oncogenic transformation, we generated *β-catenin^GOF/+^;Cdx2^f/+^;Villin-Cre^ERT^*^2^ mice (homozygous *Cdx2* loss is lethal). Following tamoxifen induction, expansion of crypt progenitors was accelerated on the *Cdx2^f/+^* background, suggesting that the fetal-like/regenerative state is important in early tumorigenesis (Fig. 6C). In conjunction with sustained *β-catenin^GOF/+^* expression, the decrease in *Cdx2* levels was sufficient to reduce expression of the ISC marker *Olfm4* and promote tissue dysplasia (Fig. S6A). RNA-seq analysis of the corresponding epithelia (control, *ß-catenin^GOF/+^* and *Cdx2^f/+^;ß-catenin^GOF/+^*) confirmed that the fetal-enriched and revival gene sets were elevated upon heterozygous *Cdx2* loss-of-function (Fig.6D), as was the YAP signaling signature characteristic of ISC regeneration (Fig. S6B). Thus, activation of a fetal-like/YAP/regenerative transcriptional program appears to facilitate the speed of oncogenic Wnt-driven transformation.

**Figure 6.**
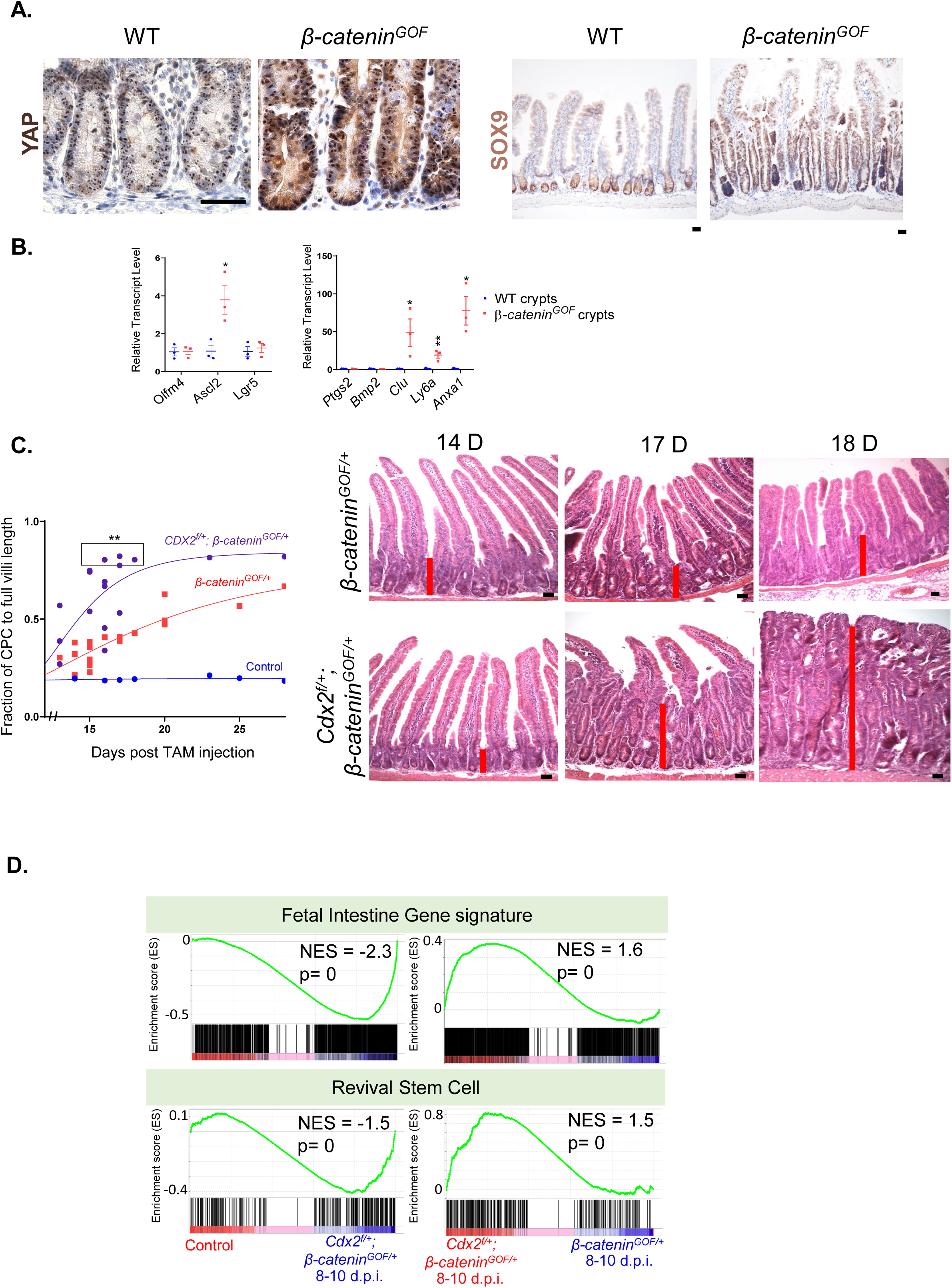
Regenerative-type growth markers are elevated in transformed tissue and loss of CDX2 leads to increased regenerative growth and faster tissue transformation in the β-catenin^GOF/+^ model. (A) Increased YAP and SOX9 immunostaining after 4 days of *β-catenin^GOF^* expression in the epithelium. Basal levels of YAP and SOX9 expression are localized to intestinal crypts in control mice (n = 3 biological replicates). Scale bar = 50 µm. (B) qRT-PCR analysis indicates a regenerative response in *β-catenin^GOF^* crypts as evidenced by upregulation of *Anxa1*, *Ascl2*, *Clu* and *Ly6a* transcripts (n = 3 biological replicates, Mann–Whitney U-test, p < 0.01** and p < 0.05*). (C) Loss of one *Cdx2* allele allows for an increased rate of tissue transformation. Rate of transformation was measured using proximal duodenum from control, *β-catenin^GOF/+^* or *Cdx2^f/+^; β-catenin^GOF/+^* mice. Unpaired t-test was performed on measurements between 15 and18 days post tamoxifen to show a difference in tissue transformation between mutants (p = 0.0051). Each point on graph represents one biological replicate. Histology (H&E) of representative samples with a red bar that highlights representative expansion of crypt-progenitor cell (CPC) zone, which was quantified as fraction of CPC to full villi length. Scale bar = 50µm. (D) GSEA analysis using bulk RNA-seq of intestinal crypts show an enrichment of fetal intestine and revival stem cell gene signatures in *β-catenin^GOF/+^* mice, which is further exacerbated by partial deletion of *Cdx2* in *Cdx2^f/+^; β-catenin^GOF/+^* mice. Intestinal crypts were collected 8-10 days after *β-catenin^GOF^* expression.

## Discussion

Stromal fibroblasts are an important source of ligands that mediate adaptation of the cellular niche. Intestinal fibroblasts have been proposed to fall under four categories of Pdgfra+ expressing cells: small intestinal crypt fibroblasts (siCF1 and siCF2), villus fibroblasts (VF) and villus tip fibroblasts (VTF).^14^ Here, we have carried out a single-cell survey of mesenchyme cell populations of mice small intestine in response to aberrant β-catenin activation in the epithelium.

Activation of the canonical WNT pathway in colorectal cancer can be mediated by loss of function of negative regulators of the pathway in epithelial cells, known as non-ligand dependent activation of the pathway, or by ligand dependent activation of the pathway, such as stromal upregulation of R-spondins.^64^ Downstream activation of canonical WNT signaling in intestinal stem cells via APC inactivation can lead to the onset of a Sox9-mediated aberrant stem cell state.^30^ In a similar manner, APC loss of function in intestinal organoids (benign adenoma model) induces an oncofetal regression in a subset of stem cells mediated by a decreased activity of the Retinoid X Receptor (RXR) and reduced chromatin accessibility by CDX2 and HNF4.^65^ In agreement with these studies, we found that activating mutations of β-catenin promotes activation of regenerative gene signatures in epithelial cells despite relatively normal differentiation potential of the mutant β-catenin epithelium. Although different crypt progenitors can revert to an earlier uncommitted state, it remains to be determined if there is a specific cell of origin for the regenerative-state in early initiation of CRC tumors. Additionally, there may be multiple modes of activating the regenerative state, for example, SOX17 can initiate a fetal program in early tumorigenesis to prevent activation of the adaptive immune system.^66^

A prolonged and sustained increase of Wnt/β-catenin signaling in the intestinal epithelium promotes a hyperplastic growth previously defined as a crypt progenitor cell (CPC) phenotype.^39,67^ Expression of a single allele of mutant β-catenin is sufficient to drive transformation of the small intestinal epithelium but not of the colon mucosa due to higher levels of E-cadherin in this compartment.^67^ E-cadherin is an important adhesion molecule that mediates cell-to-cell contacts,^68^ and loss of E-cadherin promotes a regenerative cell state during dissemination of colon cancer cells.^69^ Similar to previous studies, we found that the degree of WNT-mediated tissue transformation is highest in the proximal small intestine; thus, our analysis of regenerative-like cellular states was restricted to this region. Nonetheless, using a colon-specific Cre, we detected elevated cell proliferation in the proximal colon after 60 days of inducing mutant β-catenin expression. Further work will elucidate whether E-cadherin perturbations play a role in regulating abnormal cellular plasticity in tumor initiation.

Stromal fibroblasts are an important component of the tumor niche that can support tumor cell plasticity, which is an emergent hallmark of cancer stemness.^70^ In colorectal cancer, heterogeneous subpopulations of cancer associated fibroblasts (CAFs) can exhibit tumor-promoting or tumor-retarding properties.^71^ Multiple studies have proposed the origin of CAFs from four different sources: activation of resident fibroblasts, circulating bone marrow-derived mesenchyme stem cells, differentiation from epithelial or endothelial cells, and even transdifferentiation of pericytes.^72^ TGFB activity in CAFs is associated with poor prognosis of colorectal cancer patients. In the stromal compartment, CAFs are the main producers of TGFB, which is used to create immune-suppressive conditions in the tumor microenvironment.^73^ The investigations described here show that TGFB signaling is induced in Pdgfra+ fibroblasts in response to the initial transformation of the WNT-hyperactive epithelium. Activation of TGFB signaling is associated with expression of pro-regeneration ligands that could enhance the oncogenic WNT-induced regenerative state of the intestinal epithelium. Moreover, these fibroblasts activate inflammatory and fibrotic genetic programs that can impinge on the tumor microenvironment of CRC initiation. Mechanistically, we find that Pdgfra+ fibroblasts require TGFB activity to sustain the regenerative-type growth of wild type and β-catenin mutant enteroids. Although enteroids harboring a *β-catenin* mutation can induce the regenerative program in the absence of a stromal component, TGFB activity is required to maintain the regenerative state in culture. *β-catenin* mutant enteroids do not require intrinsic activation of the TGFB pathway to maintain the regenerative state when co-cultured with TGFB-activated Pdgfra+ fibroblasts. Taken together, we propose that TGFB activity in PDGFRA+ fibroblasts plays an important role in creating a tumor niche that favors early activation of regenerative and developmental programs in the oncogenic epithelium, and that the early activation of regenerative growth accelerates the initial stages of tumorigenesis.

## Materials and Methods

### Animals

Animal protocols and experiments were regulated by Rutgers Institutional Animal Care and Use Committee. Mice strains used in this study are listed in Table S1. Mice 6 weeks of age were treated with tamoxifen (Sigma T5648) at 50 mg/kg/day for 4 consecutive days by intraperitoneal injection, except for lineage tracing experiments in which a single of tamoxifen injection was administered at 100 mg/kg/day. Male and female mice were given equal consideration in experimental design, and littermates were used when available for matched experimental sets.

### Histology

Freshly harvested tissue was collected and fixed in 4% paraformaldehyde solution overnight at 4°C, and paraffin sections (5µm-thickness) were generated. For paraffin embedding, duodenum or jejunum tissues were dehydrated with increasing concentrations of an ethanol series and processed with xylene/paraffin before embedding. For cryo-embedding, the proximal one-third of duodenum was fixed in 4% paraformaldehyde overnight at 4°C, washed in cold PBS, and equilibrated in 20% sucrose before freezing in OCT compound (catalog no. 4583; Tissue-Tek). For immunohistochemistry, antigen retrieval was performed with 10 mM sodium citrate buffer.

Primary antibody incubation was performed overnight at 4°C in all cases. Primary antibodies used in the study are listed on Table S2. Slides were developed using 0.05% DAB and 0.015% hydrogen peroxide in 0.1 M Tris after incubation with corresponding secondary antibody and ABC-Vectastain HRP Kit (Vector Labs). Slides were counterstained with hematoxylin.

### Intestinal crypt isolation

Dissected intestine was flushed with cold PBS, cut into 0.5 cm open fragments, and incubated with 3mM EDTA/PBS at 4°C for 5 minutes, 10 minutes and 25 minutes (replacing with fresh EDTA/PBS for every step). Tissue was then manually agitated to release epithelium, which was filtered through a 70 µm cell strainer to separate crypts from villi. Crypts were pelleted by centrifugation at 200 g for 5 minutes at 4°C and rinsed with cold PBS before collection in TRIZOL for bulk RNA-seq or resuspension in Matrigel for enteroid cultures.

### Enteroid cultures

Crypts isolated from proximal duodenum were cultured in BME-R1 Matrigel (R&D, Cat no. 3433-005-R1). 80-100 crypts were plated for each 25 μl droplet of Matrigel. Enteroids were cultured in in previously established ENR medium.^47^ For treatments, enteroids were cultured with proper vehicle control or TGFB1 (Peprotech, Cat no. 100-21) at a final concentration of 2 ng/mL, BMP2 (R&D 355-BM) at a final concentration of 100 – 400 ng/mL, (16,16-dimethyl PGE_2_ (Cayman, Cat no. 14750) dissolved in ethanol at a final concentration of 10 µM, EREG (R&D Systems, Cat. no. 1068-EP-050) dissolved in 0.1% BSA at a final concentration range of 1 ng/mL – 1 µg/mL, IL-11 (Peprotech, Cat. No. 220-11) at a final concentration of 0.4 µg/mL, SB525334 (Selleckchem Catalog No. S1476) dissolved in DMSO at a final concentration of 10 µM. Enteroids were imaged using an AxioVert 200M inverted microscope (Zeiss) with a Retiga-SRV CCD (Qlmaging), or Agilent BioTek Cytation C10 Cell Imaging Multimode Reader.

### Co-culture assays and FACS cell sorting

Duodenum-derived enteroids, control or *β-catenin^GOF^*, were co-cultured with 50,000 Pdgfra-lo cells and plated in 20ul Matrigel droplets in culture medium lacking EGF, R-SPONDIN1 and NOGGIN. Culture media was replaced every day for optimal growth conditions. Pdgfra-lo controls were seeded in Matrigel without organoids. After 7 days of coculture, enteroids and Pdgfra-lo cells were physically and enzymatically separated from Matrigel using Recovery Cell Solution (Corning 354253). Then, Matrigel-free enteroid and Pdgfra-lo cells were dissociated into single cells by incubating with TrypLE (Gibco 12604013) at 37°C. Cells were then washed twice with 1% BSA/PBS and filtered with a 40-μm cell strainer. For cell sorting, cells were incubated in 1% BSA/PBS with 200 U/ml of DNase I. GFP+ cells were sorted with Beckman Coulter Astrios EQ High Speed Cell Sorter according to the described culture conditions. Dead cells were eliminated using 0.5 μg/ml DAPI. Kaluza analysis 2.1.3 software was used for FACS data analysis. For treatments, co-cultures were treated daily with SB525334 (Selleckchem Catalog No. S1476) dissolved in DMSO at a final concentration of 10 µM.

### Mesenchyme cell isolation

For single cell RNA-seq, mesenchyme cells were derived from proximal small intestine from control or *β-catenin^GOF^* mice using as previously described.^1^ For Pdgfra-lo primary monocultures or enteroid co-cultures, mesenchyme cells were isolated as previously described^44^ with minor modifications. Briefly, freshly harvested intestine was flushed with cold PBS, cut into 2 cm open fragments, followed by two incubation steps under constant rotation with pre-digestion buffer (HBSS containing 10% FBS, 10 mM HEPES and 5 mM EDTA) at 37°C for 20 minutes twice.

Next, tissues were rinsed with wash buffer (HBSS containing 2% FBS, 10 mM HEPES), and incubated with pre-warmed digestion buffer (RPMI medium containing 10% FBS, 1% P/S, 15 mM HEPES, 25 U/mL of collagenase IV (Worthington LS004186), 100 U/mL of DNase I (Sigma D4513), 0.3 g/100 mL of <u>Dispase</u> II (Gibco 17105041)) and rotated at 37°C for 30 minutes. Suspension of digested tissue was vortexed vigorously for 20 seconds every 10 minutes to release single cells. Cell solution was passed through a 40 μm cell strainer and cells were pelleted by centrifugation at 400 g for 5 minutes at 4 °C and resuspended in RPMI medium containing 10% FBS. Cells were cultured in Advanced DMEM/F12 (Gibco 12634-010) medium, containing 10% FBS (Gibco 26140-095), 1% penicillin and streptomycin (Invitrogen 15140-122), 1% HEPES (Gibco 15630-080) and 1% Glutamax (Gibco 35050-061).

### RNAscope in situ hybridization and immunofluorescence

RNAscope in situ hybridization and immunofluorescence for RNA localization were conducted using RNAscope Multiplex Fluorescent Reagent Kit v2 Assay (ACD 323110) as previously described (Holloway et al., 2021). Mice intestines were dissected and fixed as swiss rolls for 24 h in 10% normal buffered formalin at room temperature followed by a methanol series dehydration steps and rehydration with 70% ethanol before addition of paraffin. 5 μm paraffin sections were generated, baked for 1 h at 60°C, deparaffinized and pretreated with hydrogen peroxide (ACD kit) for 10 minutes, antigen retrieval (ACD 322000) for 15 minutes and protease plus treatment (ACD kit) for 30 minutes. Probe used was Mm-Ereg (437981-C2) and diluted in multiplex TSA buffer (ACD 322809). After RNAscope, sections were co-stained with mouse-anti-Ecadherin (BD Transduction Labs 6101, diluted 1:1000) and goat anti-PDGFRa (R&D systems AF1062, diluted 1:200). Secondary antibodies were donkey anti-goat IgG (Jackson ImmunoLabs 715-585-147) and donkey anti-mouse IgG (Biotium 20827) both diluted at 1:500. Slides were mounted with Prolong Gold and imaged on a Nikon AXR confocal microscope.

### RNA extraction and qRT-PCR

For enteroid monocultures, Pdgfra-lo cell monocultures, or enteroid co-cultures, Matrigel droplets containing enteroids, pdgfra-lo cells or both were homogenized in Trizol (Invitrogen 15596018), followed by RNA extraction with the QIAGEN RNeasy Micro Kit according to the manufacturer’s instructions. Freshly harvested crytps or whole tissue snips were also homogenized in Trizol an RNA was extracted following manufacturer’s instructions. For qRT-PCR, cDNA was synthesized using SuperScript III First-Strand Synthesis SuperMix (Invitrogen 18080-400). qRT-PCR analysis was conducted with customized gene-specific primers (sequences available upon request) and SYBR Green PCR Master Mix (Applied Biosystems, 4309155). The 2−ΔΔCt method was applied to calculate the fold change of relative transcript level using proper internal controls.

### Single Cell RNA-seq analysis

For each sample, data was cleansed in Cell Ranger^74^ (v6.0.0) to remove low quality reads and unrelated sequences, and data was aligned to the mouse reference genome (mm10). Cells with greater than 1 percent mitochondrial counts were removed as well in this process.

Matrices from single-cell counts were processed and analyzed through Scanpy^75^ (v1.8.2) and custom Python scripts on Google Colab. Barcodes were filtered first on Cell Ranger (v6.0.0) to include only cells with greater than 10,000 transcripts. Doublet filtering utilizing the Sam Wolock Scrublet^76^ routine was then used to remove neotypic doublets in each sample. This technique simulates doublets to create a k-nearest neighbor classifier and run a PCA to then remove likely doublets. Low complexity transcriptomes were filtered from the data by removing all barcodes with less than 100 expressed genes. Transcript UMI counts were stored in a transcripts x cell table which was total counts normalized and then log normalized. Afterwards, the top 2000 highly variable genes were selected. The data was scaled at the end to unit variance and zero mean. The single cell data was projected into a 50-dimensional principal component analysis (PCA) subspace. A (k = 20) nearest-neighbor graph was created using Euclidean distance and from the neighbor graph, clustering was performed utilizing the Leiden (Traag *et al*, 2019) community detection algorithms. Cells were represented in a uniform manifold approximation and projection plane, and cell clusters were defined via canonical marker gene composition.

### Processing of Single-Cell RNA-Seq Data with Seurat

For the goal of identifying candidate cell-cell communications using Cell Chat, Seurat was used to pre-process data. The day three mutant sample was not utilized for Seurat analysis due to a gene expression profile representing a transition state from control to mutant. Matrices from single-cell counts were processed and analyzed through Seurat^77^ (v4.0) and custom R scripts. Unique molecular identifiers (UMIs) were counted for each gene in a cell and cells were sorted by their barcodes. To remove possible doublets, cells with greater than 6,500 genes/feature counts were removed and to remove low-quality cells, cells with less than 200 genes/feature counts were removed. Gene expression measurements for each cell were then log normalized. The top 2000 highly variable genes were selected afterwards. The data was scaled so mean expression across cells was 0 and variance was constant at one. PCA was performed on the scaled data. To determine the dimensionality of the dataset and reduce technical noise, the JackStraw^78^ procedure was utilized to construct a ‘null distribution’ of feature scores to identify principal components with the strongest enrichment (ones with low p-values). An elbow plot was run alongside this to confirm the number of dimensions. Two analysis runs of Seurat were performed, for control and mutant samples. A k-nearest neighbor graph was created based on the Euclidean distance in the PCA dimensions (23 for mutant, 29 for control) of the data. Clustering was then performed utilizing the Louvain^79^ algorithm. Cells were projected into a two-dimensional tSNE plane, and cell clusters were found through their marker gene composition. Gene expression data matrix and cell labels for each sample created by Seurat were loaded into CellChat^56^ (v1.1.3) to create a CellChat object. The CellChatDB mouse literature-supported ligand-receptor interactions database was utilized in the analysis which consisted of 59.9% paracrine/autocrine signaling interactions, 21.4% extracellular matrix (ECM)-receptor interactions, and 18.7% cell-cell contact interactions. The data was preprocessed to infer cell state-specific communications through identification of overly expressed ligands or receptors in cell groups and reciprocal ligand-receptor over expression.

Inference of cell-cell communication networks was then performed by giving each interaction in the preprocessed data a probability value and then modeling the probabilities utilizing gene expression in combination with prior known ligand-receptor interactions using the law of mass action. Communication probability on a signaling pathway level is then found by amalgamating all the probabilities of ligand-receptor interactions within each pathway.

### Quantification and statistical analysis

The data is presented as mean ± SEM, and statistical comparisons were generated using one-way ANOVA followed by Tukey’s multiple comparisons test with the GraphPad Prism version 8.0.2, Student’s t-test or Mann–Whitney U-test at p < 0.0001^∗∗∗∗^, p < 0.001^∗∗∗^ p < 0.01^∗∗^ or p < 0.05^∗^.

## Acknowledgements

M.P.V. is supported by grants from the National Institutes of Health (NIH R01 CA190558). M.P.V. and J.R.S. are supported by the Intestinal Stem Cell Consortium from the National Institute of Diabetes and Digestive and Kidney Diseases (NIDDK) and National Institute of Allergy and Infectious Diseases (NIAID) of the NIH under grant number U01 DK103141. K.D.W. is supported by a grant from the NIH (R01 DK121166). The authors acknowledge the Office of Advanced Research Computing (OARC) at Rutgers University for providing access to the Amarel cluster and associated research computing resources that have contributed to the results reported here. The research was also supported by flow cytometry/cell sorting core facility at Environmental and Occupational Health Sciences Institute (EOHSI) at Rutgers University. This work benefited from the Cancer Center Support Grant (CCSG, P30CA072720) from the National Cancer Institute.

## Author Contributions

O.P.C. conceived and designed the study, performed benchwork, sequencing data processing and bioinformatics, collected and analyzed the data; and wrote the manuscript. O.P., A.W., R.K., M.A., A.L., S.W. and K.T. contributed to benchwork, P.R., E.F., I.P., K.D.W., and J.P. contributed to benchwork and data collection. E.M., D.E.W., P.H., D.F. and H.S. contributed to sequencing data processing and bioinformatics, J.R.S., N.J.B. and R.A.S provided experimental materials or instruments. M.P.V conceived, supervised the study and wrote the manuscript.

---

**Supplementary Figure 1.**
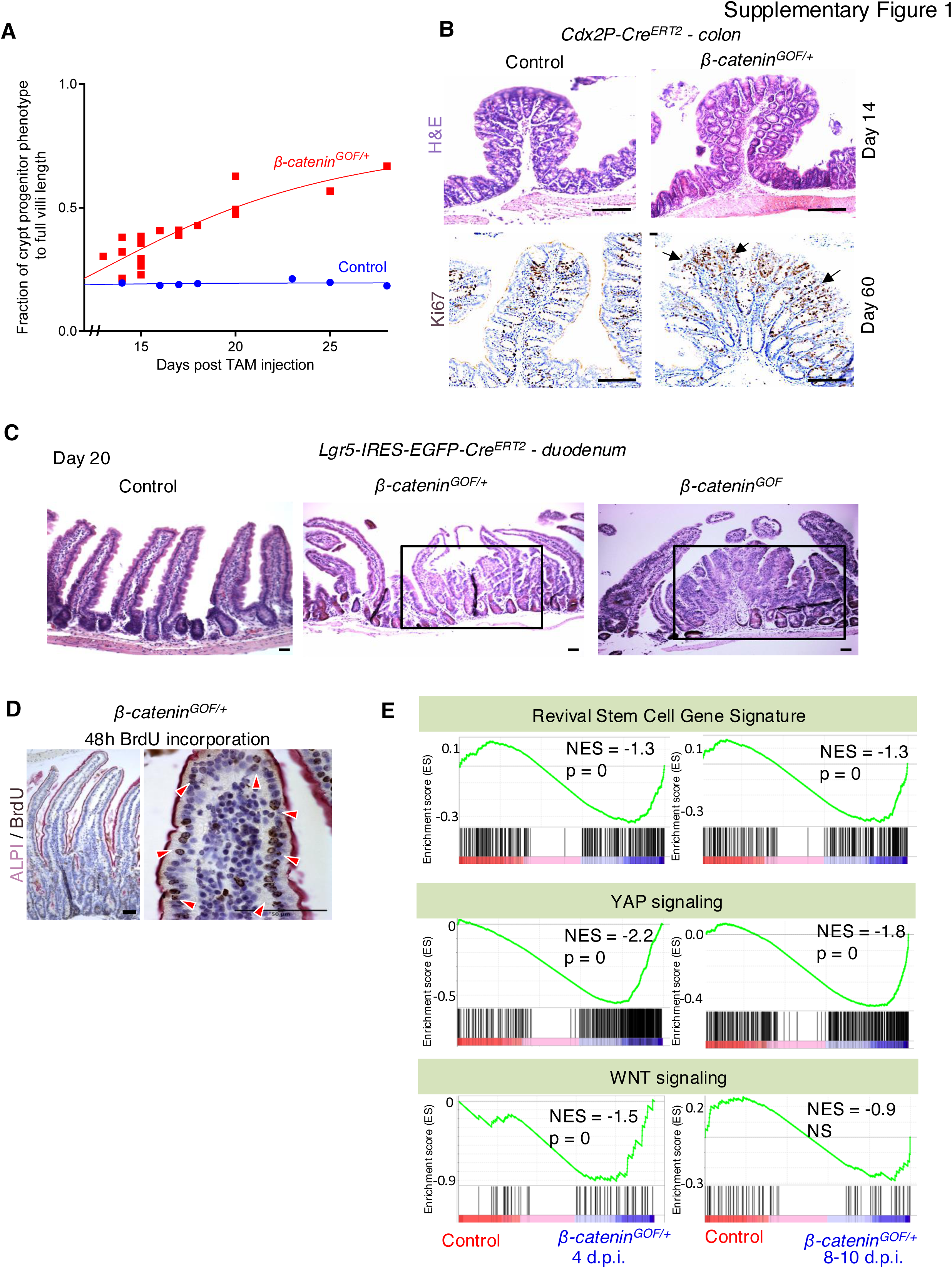
(A) Quantitation of H&E analysis to compare magnitude of crypt-progenitor cell phenotype (CPC) in proximal duodenum between control and *β-catenin^GOF/+^; Villin-Cre^ERT^*^2^ mice. (B) Histology (H&E) of control and *β-catenin^GOF/+^; Cdx2P-Cre^ERT^*^2^ mice indicate no difference in transformation of proximal colon at day 14. Immunohistochemistry of Ki67 show increased cell proliferation in *β-catenin^GOF/+^* colonic tissue at day 60 (n = 3 biological replicates), consistent with the latency in tissue transformation observed in the small intestine. (C) H&E images showing that heterozygous or homozygous ß-catenin activation in Lgr5^+^ stem cells lead to moderate tissue transformation (black box) at day 20 (n = 3 biological replicates). (D) Pulse labeling of *β-catenin* mutant stem cells with BrdU (48h pre-harvest) at day 20 after expression of the mutant *β-catenin^GOF^* allele in *β-catenin^GOF/+^; Villin-Cre^ERT^*^2^ mice. BrdU^+^ cells in villi also exhibit staining of differentiation marker alkaline phosphatase (AP) (n = 3 biological replicates) (E) RNA-seq analysis of primary crypt epithelium shows activation of revival stem cell, YAP signaling, and WNT signaling gene signatures in control vs *β-catenin^GOF/+^* crypt cells at day 4, or day 8-10 post-induction of a single *β-catenin^GOF^* mutant allele using the *Villin-Cre^ERT^*^2^ driver. *Scale bars: 50 µm.

**Supplementary Figure 2.**
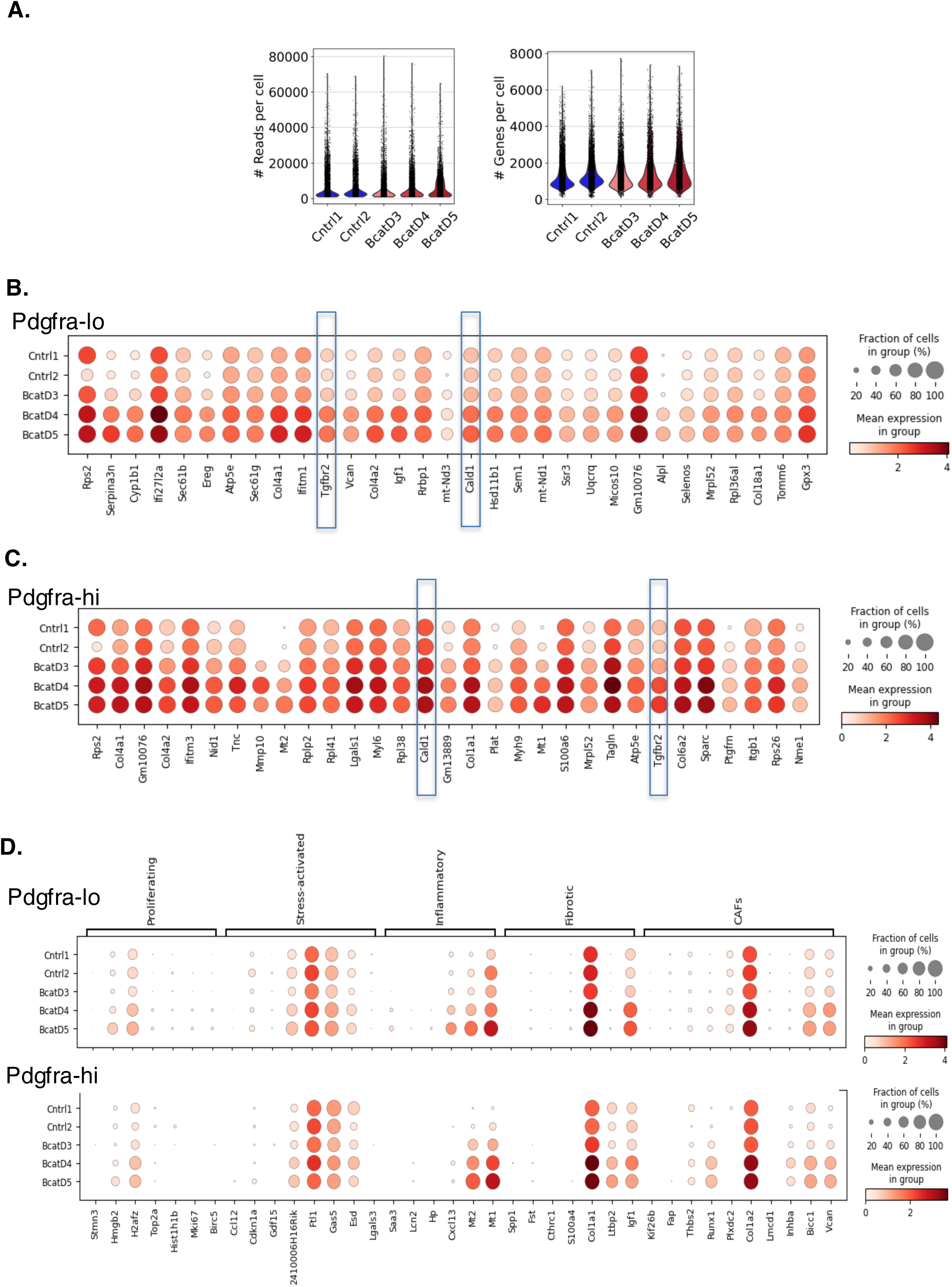
(A) Violin plots of total number of reads and genes detected in each scRNA-seq sample. (B) scRNA-seq dot plots of top 30 upregulated genes in Pdgfra-lo cells in control and *ß-catenin^GOF^* mice. TGFB pathway-related genes *Tgfbr2* and *Cald1* are elevated with higher WNT activity in epithelium. (C) scRNA-seq dot plots of top 30 upregulated genes in Pdgfra-hi cells in control and *ß-catenin^GOF^* mice. TGFB pathway-related genes *Tgfbr2* and *Cald1* are elevated with higher WNT activity in epithelium. (D) scRNA-seq analysis indicate that markers for proliferating, inflammatory, fibrotic and CAFs fibroblasts are elevated in Pdgfra-lo and Pdgfra-hi cells in response to WNT hyperactivation in the epithelium.

**Supplementary Figure 3.**
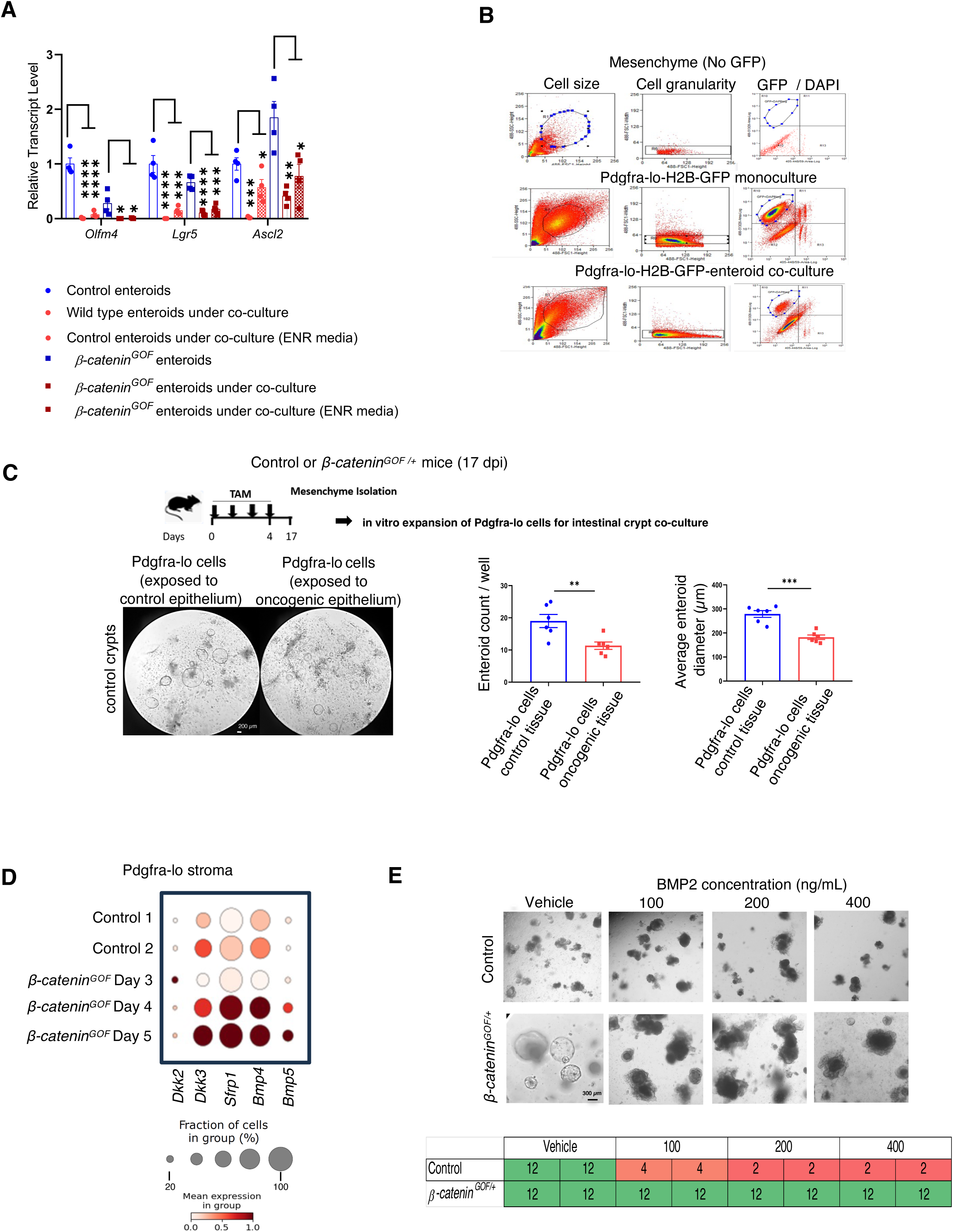
(A) qRT-PCR analysis indicates that Lgr5+ stem cell markers are downregulated in both control or *β-catenin^GOF^* enteroids in co-culture with Pdgfra-lo cells (n = 4 biological replicates, one way ANOVA with Tukey post hoc test, p < 0.0001****, p < 0.01** and p < 0.05*). (B) FACS sorting strategy to isolate Pdgfra-H2BeGFP cells from enteroid co-cultures. (C) Pdgfra-lo cells exposed to *β-catenin^GOF/+^* epithelium for 17 days does not support growth of control crypts as well as Pdgfra-lo cells from control mice. (n = 3 biological replicates, Student’s t-test at p < 0.001*** and p < 0.01**). Scale bar = 200 µm. (D) Dotplot of scRNA-seq profiles of Pdgfra-lo cells indicates an upregulation of pro-differentiation ligands in response to high WNT levels in the epithelium. (E) Extraphysiological levels of BMP2 promotes differentiation of enteroid cultures derived from *β-catenin^GOF/+^* mice and cell death in organoids derived from control mice. Table indicates viability (in days) of enteroids treated with BMP2. (n = 2 biological replicates, 3 technical replicates). Scale bar = 300 µm.

**Supplementary Figure 4.**
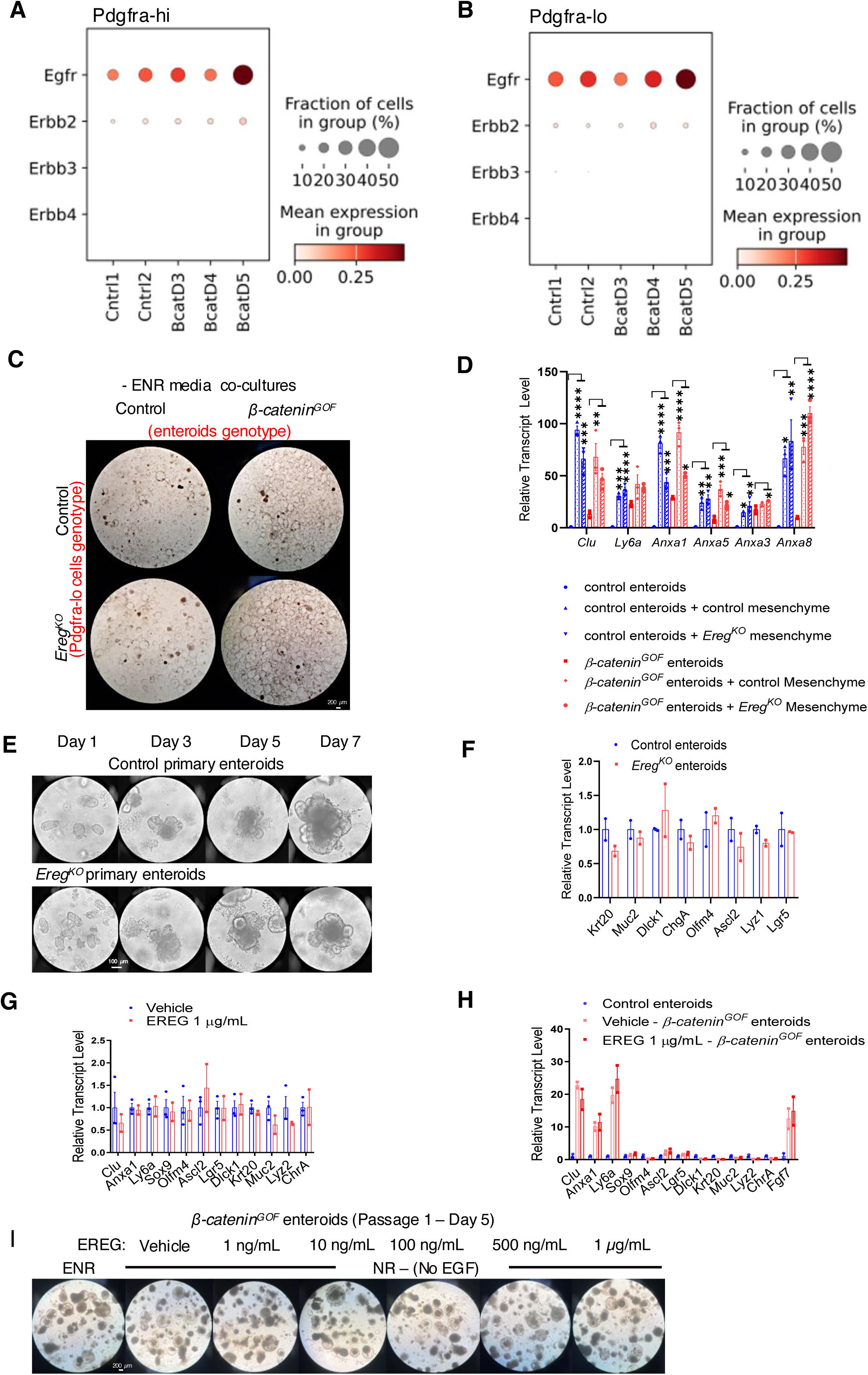
(A-B) scRNA-seq dot plots indicate highest expression of *Egfr* in both PDGFR-hi and Pdgfra-lo cells at day 5 of homoyzygous *β-catenin^GOF^* activation in the epithelium. (C) Pdgfra-lo cells derived from *Ereg^KO^* mice can sustain growth of both control and *β-catenin^GOF^* enteroids in co-culture assays (representative images of 3 biological replicates). Scale bar = 200 µm (D) Pdgfra-lo cells derived from *Ereg^KO^* mice induce similar expression of fetal/regenerative markers in both control and *β-catenin^GOF^* enteroids in co-culture assays (n = 3 biological replicates, one way ANOVA with Tukey post hoc test, p < 0.0001****, p < 0.001***, p < 0.01** and p < 0.05*). (E) Intestinal crypts derived from *Ereg^KO^* mice develop into enteroids with normal morphology resembling littermate-derived control enteroids (n = 2 biological replicates, 3 technical replicates each). Scale bar = 100 µm. (F) qPCR analysis shows that multilineage differentiation is uncompromised in *Ereg^KO^* enteroids (n = 2 biological replicates, 3 technical replicates each). (G-H) qPCR data indicates that treatment of control or *β-catenin^GOF^* enteroids with recombinant EREG (1 µg/mL) does not affect expression of a myriad of genes related to differentiation, fetal-like/regenerative, or stem cell state (n = 2 biological replicates per group, 3 technical replicates each). (I) Normal growth of *β-catenin^GOF^* enteroids cultured in an EGF-depleted medium but supplemented with different concentrations of recombinant EREG (1 ng/mL – 1 µg/mL) (representative images of 3 biological replicates). Scale bar = 200 µm.

**Supplementary Figure 5.**
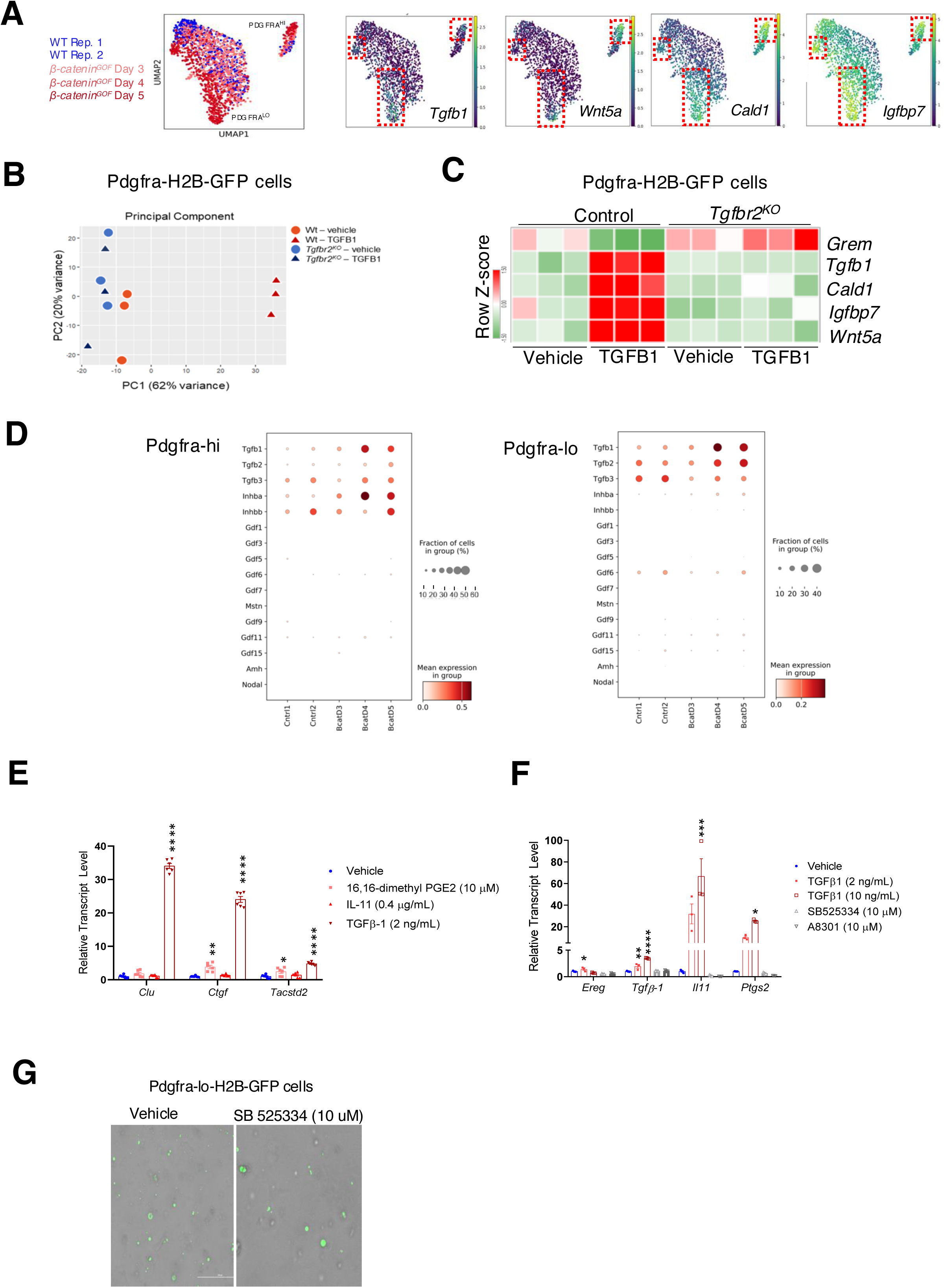
(A) UMAP of Pdgfra-lo cells show that cells exposed to hyperactive WNT epithelium have elevated expression of TGFB1 targets. Red boxes highlight portions of the clusters enriched with cells derived from *ß-catenin^GOF^* mutant mice. (B) PCA plot of control or *Tgfbr2^KO^* Pdgfra-lo cells along with its biological replicates based on their treatment with vehicle or TGFB1. (C) Heatmap of key TGFB1 targets identified by RNA-seq analysis of control or *Tgfbr2^KO^* Pdgfra-lo cells after TGFB1 treatment. (D) scRNA-seq dotplots show increased expression of *Tgfb1* and *Inhba* both Pdgfra-lo and Pdgfra-hi cells after induction of ß-catenin activity in the epithelium (E) qRT-PCR data shows that the TGFB1 ligand is the strongest inducer of fetal/regenerative genes. Among other regenerative ligands, TGFB1 promoted significant developmental reprogramming of enteroids under 24 hours (n = 6 biological replicates) (F) qRT-PCR analysis shows that induction of pro-regeneration ligands by TGFB1 in Pdgfra-lo cells can be inhibited by small molecules inhibitors, SB525334 (10 µM) and A8301 (10 µM), targeting the TGFB pathway. (G) Images showing viability of Pdgfra-lo cells after treatment with SB525334 (10 µM) (n = 3 biological replicates) Scale bars = 300 µm.

**Supplementary Figure 6.**
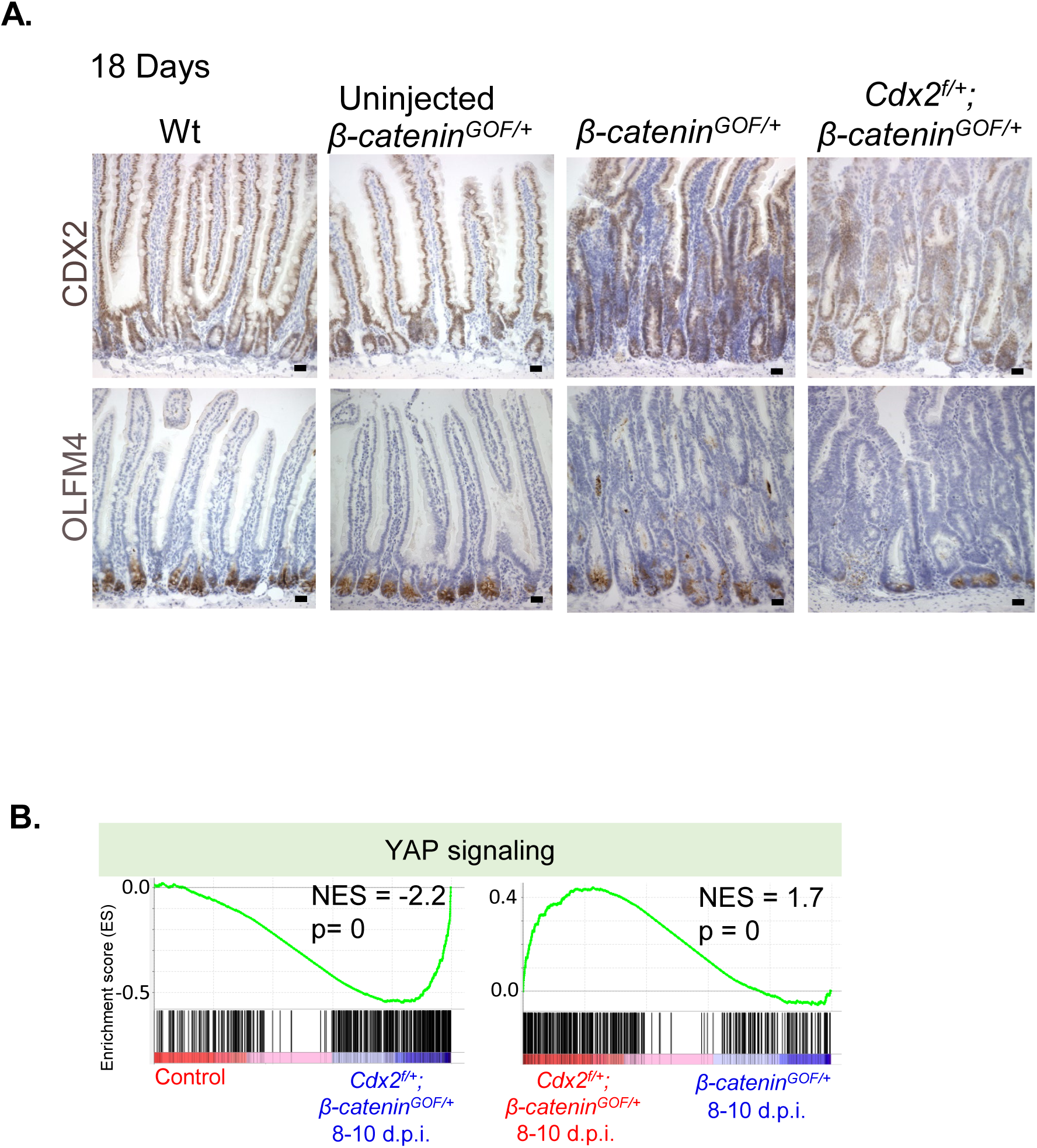
(A) Immunohistochemistry (IHC) of CDX2 and OLFM4 shows reduced CDX2 and OLFM4 levels in transformed tissue (day 18) from *Cdx2^f/+^; β-catenin^GOF/+^* mice. Scale bar = 50 µm. (B) GSEA analysis indicate that YAP activation is elevated in *Cdx2^f/+^; β-catenin^GOF/+^* epithelium. Partial inactivation of CDX2 further increases YAP activation induced by *β-catenin^GOF/+^* expression.

**Table S1.**
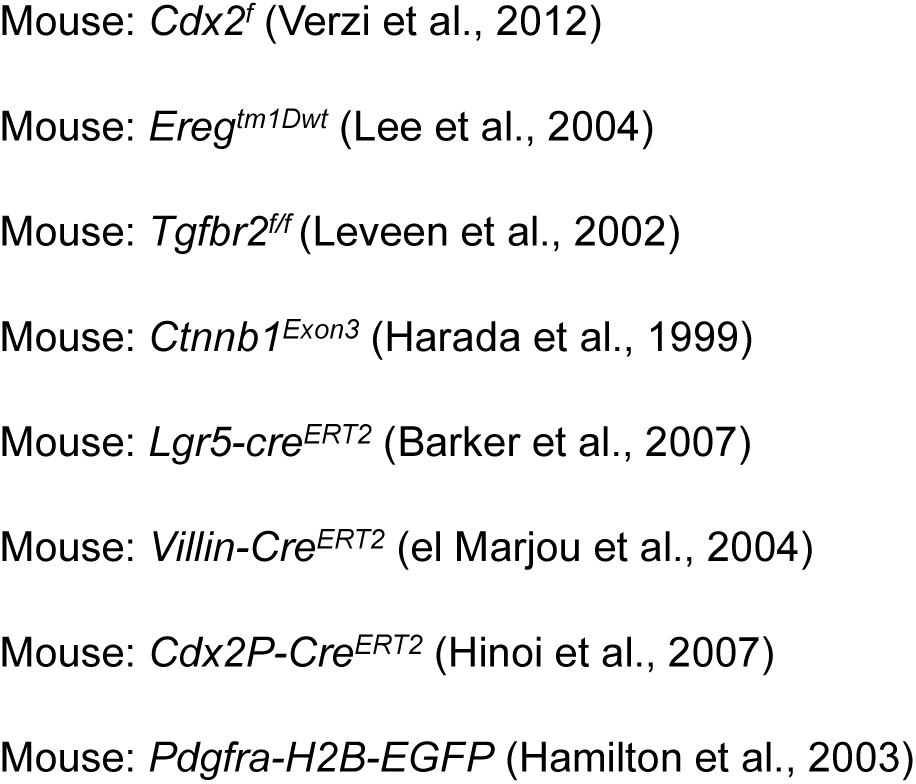
Allele References.

**Table S2.**
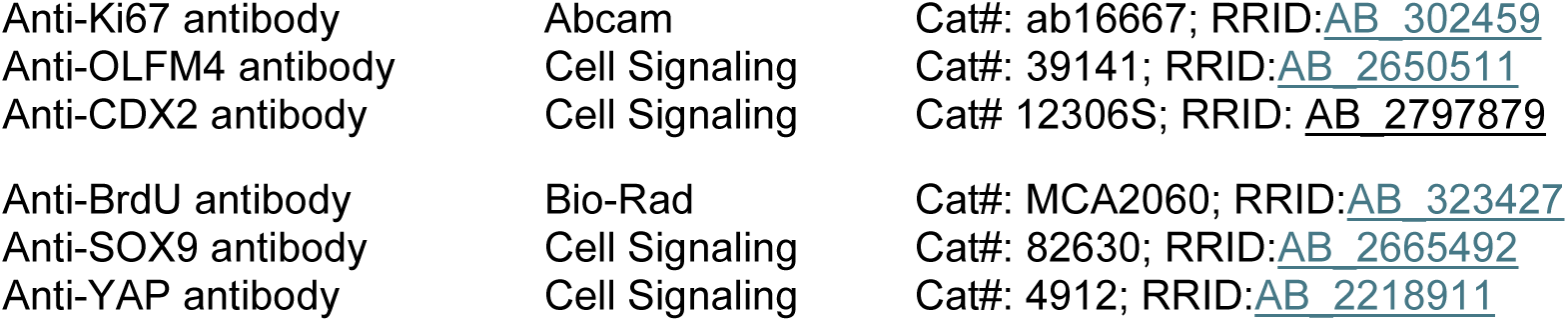
Antibodies.

|  |  |  |
| --- | --- | --- |
| Anti-Ki67 antibody | Abcam | Cat#: ab16667; RRID:AB_302459 |
| Anti-OLFM4 antibody | Cell Signaling | Cat#: 39141; RRID:AB_2650511 |
| Anti-CDX2 antibody | Cell Signaling | Cat# 12306S; RRID: AB_2797879 |
| Anti-BrdU antibody | Bio-Rad | Cat#: MCA2060; RRID:AB_323427 |
| Anti-SOX9 antibody | Cell Signaling | Cat#: 82630; RRID:AB_2665492 |
| Anti-YAP antibody | Cell Signaling | Cat#: 4912; RRID:AB_2218911 |

